# The evolution of multiply substituted isotopologues of methane during microbial aerobic oxidation

**DOI:** 10.1101/2023.11.02.565373

**Authors:** Jiawen Li, Beverly K. Chiu, Alison M. Piasecki, Xiahong Feng, Joshua D. Landis, Sarah Marcum, Edward D. Young, William D. Leavitt

## Abstract

Aerobic methane oxidation (AeOM) is an important biological sink of methane on Earth. Stable isotopes are critical tools in tracking the sources and sinks of Earth’s surface methane budget. However, the major factors that influence the two multiply-substituted (clumped) isotope signatures of AeOM, Δ^13^CH_3_D and Δ^12^CH_2_D_2_, are not well known. Here we quantify the influence of kinetics as a function of temperature, and different methane monooxygenase (MMO) enzymes as a function of copper, on the isotopologue concentrations of residual methane by the obligate aerobic methanotroph, *Methylococcus capsulatus* (Bath). We observe deviations from traditional closed-system distillation (Rayleigh) fractionation during exponential growth at high oxidation rates. We model this as a reservoir effect controlled by the ratio of oxidation rate in the cells to transport rate of methane into the cells, where environmental temperature affects both rates. We also test whether clumped isotope fractionation values vary for the particulate versus soluble MMOs, but the results show minimal differences. We further determine that the back reaction (re-equilibration) of methane with medium water is unlikely. Together, the observations and model demonstrate that at low oxidation-to-transport ratios, the clumped isotope signatures follow canonical Rayleigh fractionation, whereas at high ratios, more positive Δ^12^CH_2_D_2_ values result, deviating from simple Rayleigh-like trajectories. This study shows that the methane oxidation-to-transport ratio is a critical influence on clumped isotope signatures of AeOM that should be considered when interpreting the isotopic data of natural methane samples in both open and closed systems.

## 1. Introduction

Methane is a powerful greenhouse gas that has played a role in Earth’s climate for billions of years (Catling et al., 2001; Frieling et al., 2016). Since the Industrial Revolution, methane has accumulated in Earth’s atmosphere faster and to a greater extant than any time in at least the last 800 kyr (Ruppel and Kessler, 2017), with significant consequences predicted for extant and future climate (Dlugokencky et al., 2011; Saunois et al., 2020). Methane is also a major component of natural gas, powering vast swaths of the human economy (Hamak and Sigler, 1991; Faramawy et al., 2016). Beyond Earth, the presence and isotopic compositions of methane on other planetary bodies may enable us to differentiate biological from non-biological processes (McKay and Smith, 2005; McKay et al., 2008; Yung et al., 2018; Thompson et al., 2022).

The major sources of methane on Earth include those that are either natural or anthropogenic, as well as biotic or abiotic in origin (Dlugokencky et al., 2011; Kirschke et al., 2013; Saunois et al., 2020). Anthropogenic methane sources include release during fossil fuel extraction, biomass burning, coal mining, agriculture, and waste disposal (i.e. anoxic microbial activity). Natural sources include the microbial degradation of organic matter (e.g. in anoxic wetlands, lakes, oceans, and ruminant guts) and the release of methane from permafrost and gas hydrates. Abiotic methane synthesis is the product of water-rock reactions in hydrothermal systems, including, at lower temperatures, serpentinization (McCollom and Seewald, 2001; Etiope and Sherwood Lollar, 2013). The major methane sinks on Earth include abiotic oxidation in the tropo- and stratosphere due to interactions with OH and Cl radicals (Ehhalt, 1974; Cicerone and Oremland, 1988), as well as microbial oxidation, a.k.a “methanotrophy” (Curry, 2007). Microbial methanotrophy can be further categorized into the anaerobic oxidation of methane (AOM) and the aerobic oxidation of methane (AeOM). Among them, AeOM plays an important role in methane consumption in peat soils (Oremland and Culbertson, 1992), ocean water (Qin et al., 2022; Mao et al., 2022), and in the atmosphere (Greening and Grinter, 2022). However, there are large discrepancies between top-down and bottom-up estimates of Earths’ methane budget (Saunois et al., 2020). In part, this is due to a lack of reliable tools to track the sources and sinks of methane via different pathways.

Methane made and destroyed by abiotic and biological processes is possessed of unique isotopic information that may provide means to identify and track the sources and sinks of methane (Schoell, 1988; Whiticar, 1999; Douglas et al., 2017; Young et al., 2019). Several studies investigated bulk isotope (*δ*^13^C and *δ*D) fractionation of methane oxidation by lab-cultured aerobic methanotrophic bacteria (Coleman et al., 1981; Templeton et al., 2006), as well as in marine and fresh waters (Kawagucci et al., 2021; Thottathil et al., 2022). Technological advances in the measurement of doubly substituted methane isotopologues (^13^CH_3_D and ^12^CH_2_D_2_, a.k.a. clumped isotopologues) have allowed for development of a novel tool for tracking the formation and consumption of methane gas (Stolper et al., 2014; Wang et al., 2015; Young et al., 2017). At thermodynamic equilibrium, the distribution of ^13^CH_3_D and ^12^CH_2_D_2_ within a population of methane molecules correlates with temperature (Cao and Liu, 2012; Liu and Liu, 2016; Eldridge et al., 2019). Therefore, at equilibrium, methane clumped isotope values can be applied as thermometers for tracking the formation temperature (Stolper et al., 2014; Young et al., 2017; Thiagarajan et al., 2020). In contrast, microbial methane often shows disequilibrium clumped isotope values (Stolper et al., 2015; Wang et al., 2015; Douglas et al., 2016; Young et al., 2017; Gruen et al., 2018; Giunta et al., 2019; Taenzer et al., 2020), distinct from those of abiotic or high-temperature equilibrated methane (Young et al., 2017; Gonzalez et al., 2019; Labidi et al., 2020). These disequilibrium values allow us to distinguish methane gases from different sources. However, one of the major problems hindering the application of methane clumped isotope values in natural systems is that the original clumped isotope compositions of a given methane source are susceptible to alteration by microbial methane oxidation, both aerobic (AeOM) and anaerobic (AOM) (Haghnegahdar et al., 2017; Ash et al., 2019; Ono et al., 2021; Giunta et al., 2022).

Prior studies suggest AOM drives methane clumped isotope signatures towards equilibrium at low reaction rates (Ash et al., 2019; Giunta et al., 2019; Ono et al., 2021; Zhang et al., 2021; Liu et al., 2023). In contrast, the studies to-date show AeOM generates different clumped isotope signatures from AOM (Wang et al., 2016; Krause et al., 2022; Giunta et al., 2022). However, the major controls on AeOM methane clumping remain poorly understood. To date, there are only two lab studies on the role of AeOM in setting methane clumped isotope compositions (Wang et al., 2016; Krause et al., 2022). While both are foundational in providing the first measures of AeOM clumped isotope fractionation in one (Wang et al., 2016) and two (Krause et al., 2022) clumping dimensions, neither addresses the influence of growth rate, nor enzyme-specific isotope fractionation, on methane clumped isotope compositions. Fortunately, the core metabolic processes of AeOM are well understood, and microbial strains such as *Methylococcus capsulatus* strain Bath (referred to hereafter as *M. capsulatus*) are well characterized genetically and physiologically (Hanson and Hanson, 1996), which makes them good targets for the study of the isotope fractionation during AeOM.

Given the known influence reaction rate has on the expressed isotope effects during AOM, we predict rate has an impact on the isotope fractionation of AeOM as well. Metal availability, in particular copper (Cu), controls the microbial expression of two types of methane monooxygenases (MMOs) (Stanley et al., 1983; Prior and Dalton, 1985), each of which catalyzes the first critical step in AeOM. Between the two distinct MMOs, a soluble form that uses a diiron active site (sMMO) and a membrane-bound form with a catalytic copper center (pMMO) (Ross and Rosenzweig, 2017), it remains unknown whether each type has different or similar clumped isotope fractionation. In this study, we set out to investigate the trajectory of methane clumped isotope signatures as a function of growth rates in response to growth temperature, as well as the activities of MMOs controlled by copper concentrations. We evaluate our experimental findings with a cellular fractionation model that seeks to capture the effects of transport limitation and the closed-system dynamics of the experiments. We then discuss the implications of the experimental isotope fractionation on methane in natural systems.

## 2. Method

### 2.1. Microbial culturing

All experiments in this study were performed with *Methylococcus capsulatus* (Bath) acquired from ATCC (ATCC 33009). *M. capsulatus* is an obligate aerobic methanotroph, representative of the wider diversity of aerobic methanotrophic bacteria (Bowman, 2014) that utilize methane for both carbon and energy (Hanson and Hanson, 1996). We performed two types of batch experiments to test the effect of either the two different MMOs (controlled by copper concentration) or the growth rate (controlled by growth temperature) on clumped isotope fractionations. Axenic cultures of *Methylococcus capsulatus* were grown in nitrate mineral salts (NMS) media following established methods (Wang et al., 2016; Welander and Summons, 2012). Briefly, 30 mL NMS medium was dispensed into 160-mL serum bottles, sealed with ambient air at atmospheric pressure and temperature, then injected with 20 mL of filter-sterilized ultra high purity methane (UHP, AirGas, ME). Each serum bottle was inoculated with 1 mL of starter culture at optical densities of ∼ 0.7 (OD_600_). For the differential temperature experiments, cultures were incubated at 21, 27, or 37 °C in NMS media supplemented with 5 *μ*M CuSO_4_. For the copper concentration experiments, we used basal NMS medium with starting CuSO_4_ concentrations at 0 and 50 *μ*M in addition to the 5 *μ*M used in the differential temperature experiments, with all incubations conducted at 37 °C. To test the possibility of hydrogen exchange between methane and medium water, two sets of incubations were done with NMS media made from deuterium-spiked (D-spiked) water at +1500 and +3000 ‰ *δ*D under 37 °C and with 5 *μ*M CuSO_4_ added. For simplicity, the differential temperature experiments are referred to as “37 °C”, “27 °C” and “21 °C” experiments, while the differential copper experiments are referred to as “no-Cu” (0 *μ*M) and “high-Cu” (50 *μ*M), respectively. To ensure the exchange of headspace methane with the medium liquid, bottles were shaken at 200 rotations per minute (RPM) in temperature controlled rotary shakers (Innova42, Eppendorf).

A minimum of four time points were sampled from each experiment, with five replicates per time point. The time points for sampling were determined based on the amount of residual methane in the headspace. We selected the time points so that the residual methane decrease by 10-20 % between the two adjacent time points. Typically a minimum of 60 to 80 *μ*mol of methane is targeted for isotopic measurements. Later timepoints have less, but still measurable methane with the cost of lower analytical precision. In addition, one negative control bottle (medium without inoculum) was incubated for each temperature or copper series. From the five replicate bottles, two were designated for monitoring growth (OD_600_), and the other three for methane sampling at the time of sacrifice. To minimize the chance of methane leakage, the septa of the three methane sample bottles were only punctured to add the methane, the cells, and to measure residual methane at the time of sacrifice. Microbial growth was quantified by changes in OD_600_ over time. The growth rates and cell doubling times were calculated based on the optical density in the exponential growth phase, following the equations in Brock et al. (2003):

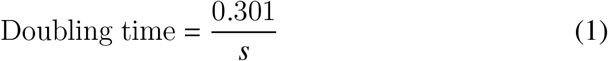

where *s* is the slope of a linear function from the plot of log_10_(OD) over time during the exponential phase (Figure B.9). The errors in doubling time were propagated from the errors of the linear regressions. At each time point of sampling, all five replicates were sampled for residual methane amount by gas chromatography (GC). The five biological replicates were then terminated by the addition of 1 mL 1M HCl to the 30-mL media, and stored inverted at 4 °C until methane extraction. Copper concentrations were measured on the two monitor bottles at the end of growth experiments. Following GC measurements, the media in the monitor bottles were centrifuged without acidification, filtered, then prepared for ICP-OES (see below).

### 2.2. Analytical techniques

#### 2.2.1. Methane concentration

Methane concentrations in the headspace were monitored by gas chromatography fitted with a flame ionization detector (GC-FID, SRI 8610C) with Grade 5.0 helium (UHP, AirGas). During the measurement, the column temperature was held at 120 °C, and the retention time of methane was consistently ∼ 1 min. A daily calibration with 1% (v/v) methane/helium standard (AirGas) was done to ensure the analytical linearity of the instrument prior to the measurements. Headspace gas was sampled from each bottle (40 *μ*L) using a calibrated gas-tight syringe with a valve (100-*μ*L; Hamilton Company) for the quantification of methane concentrations. The results were then converted to total headspace methane in each bottle. The uncertainty of each sample was propagated from the error in the GC calibration. Dissolved methane in the media was calculated following Yamamoto et al. (1976).

#### 2.2.2. Dissolved copper concentration

Dissolved copper concentrations in the medium were measured using an inductively coupled plasma - optical emission spectrometer (ICP-OES; axial-view Spectro ARCOS), and calculated from the variance-weighted average of the two emission lines, 324 and 327 nm (Landis et al., 2021). The standards were matrix-matched to the samples, with an independent calibration verification standard showing residual standard deviation typically smaller than 2%. The standards were measured every five samples to correct for the time-dependent drift. The instrument detection limit (IDL) was calculated as 3x standard deviation of blank measurements (in ng/mL). For copper, the IDL is 3.9 ng/mL (0.0614 *μ*M), which was sufficient for monitoring the change of copper concentration during the experiment. Each sample was measured twice and the average values of the two measurements were taken as the final results.

#### 2.2.3. Isotopic measurements

Prior to isotopic measurements, methane was purified from other headspace gases on a vacuum line. The procedure was modified based on the method described in Taenzer et al. (2020). The gas was introduced from serum bottles into the vacuum line through an SGE on/off valve (SMOV) with a needle attached to it. The gas was then equilibrated between the bottle and the vacuum line for 5 minutes to avoid fractionation, after which the SGE valve was closed. Water vapor and carbon dioxide were trapped in a U-trap by pure liquid nitrogen (LN_2_) at −185 °C. Methane was trapped in a U-trap containing silica gel at LN_2_ temperature, followed by entrainment in 5.5 grade He and transport to a GC (SRI GC-TCD 8610). Purification of CH_4_ from non-condensible atmospheric gases (Ar, O_2_, N_2_) and higher-order organics was accomplished using two in-series packed GC columns, one filled with 5A molecular sieve and the other with HayeSep D polymer. After GC purification, methane gas was trapped in a second U-trap filled with silica gel at LN_2_ temperature, followed by pumping away the He carrier gas. The purified methane gas was then collected in a sample vial with silica gel at LN_2_ temperature and transferred via a small-volume cold finger, also containing silica gel, to the mass spectrometer for isotopic measurements.

The isotopic compositions of headspace methane were measured on the prototype Panorama (Nu Instruments, AMETEK Inc.) high-resolution, double-focusing mass spectrometer at UCLA. To resolve two mass-18 isotopologues of methane (^13^CH_3_D and ^12^CH_2_D_2_), as well as interfering species, the instrument was run at a mass resolving power of ∼ 40,000 or above (Young et al., 2016). We utilized a combination of Faraday collectors with 10^11^Ω resistors and a single channel of ion counting for these measurements. Isotopic ratios of the samples were measured relative to a reference methane gas under two magnet current settings. The first setting placed ^13^CH_3_D ^+^ in the axial electron multiplier collector. ^13^CH_4_ ^+^/^12^CH_4_ ^+^ and ^13^CH_3_D ^+/12^CH_4_ ^+^ ratios were obtained after 6 to 20 blocks of twenty 30-sec integration cycles. The second setting placed ^12^CH_2_D_2_ ^+^ in the axial collector where ^12^CH_3_D ^+/12^CH_4_ ^+^ and ^12^CH_2_D_2_ ^+/12^CH_4_ ^+^ ratios were obtained after 16 to 40 blocks of twenty 30 sec integration cycles (Young et al., 2017). The number of blocks used for each measurement depended on the size of the methane sample, with a maximum number of 40. Fewer blocks of measurements generally resulted in larger internal 1*σ* uncertainties. The internal 1*σ* uncertainties (propagation of standard errors in the means for all blocks) of the samples are within ± 0.28 ‰ for *δ*^13^C, ± 0.09 ‰ for *δ*D, ± 0.74 ‰ for Δ^13^CH_3_D, and ± 0.86 ‰ for Δ^12^CH_2_D_2_ (Table 1).

**Table 1:**
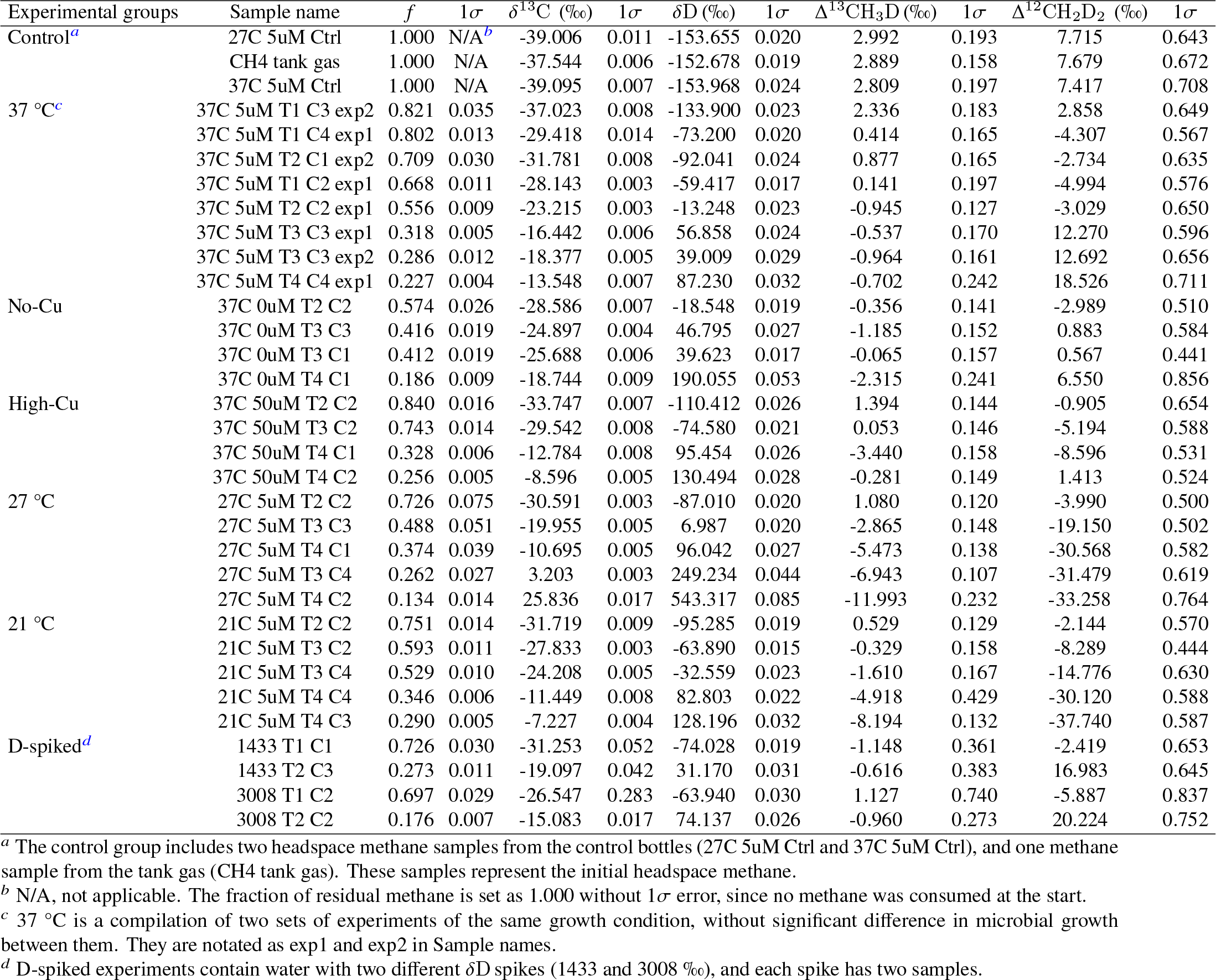
Isotopic data of headspace methane in aerobic methane oxidation by *Methylococcus capsulatus* (Bath). Experimental conditions are represented as either growth temperature or copper concentration in the column “Experimental groups”. The differential temperature experiments are conducted with 5 *μ*M Cu in the media, and the differential copper experiments are conducted under 37 °C. The 1*σ* errors are listed following each measured variables.

Isotopic compositions of medium water were measured in the Stable Isotope Laboratory at Dartmouth College following the procedures described previously by Kopec et al. (2019) using an H-Device coupled to a Thermo Delta Plus XL dual-inlet isotope ratio mass spectrometer (IRMS). Measured values were converted to the water isotope equivalent by calibration with known standards. Deuterium-spiked water samples outside the range of working standards were diluted with water of known isotopic compositions before measurement following the procedure in Taenzer et al. (2020). Uncertainties in mass spectrometry measurements and dilutions were propagated into the result of each measurement. The average values and the standard deviations among the four biological replicates in each set of experiment were taken as the final results. Isotope ratios are reported as *δ*D in ‰ relative to VSMOW, as shown in Table C.5.

### 2.3. Isotope notations and calculations

#### 2.3.1. Isotope notations

Bulk isotope ratios for methane in this study are reported using standard delta-notation (in ‰ relative to international standards):

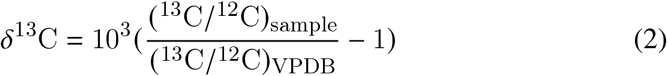

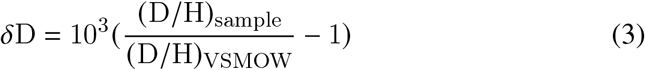

where VPDB refers to the Vienna Pee Dee Belemnite and VSMOW refers to Vienna Standard Mean Oceanic Water.

Clumped isotope compositions are expressed using the capital delta notation in ‰ relative to the corresponding stochastic distribution for the sample in which isotopes are randomly distributed among the CH_4_ isotopologues (Wang et al., 2004):

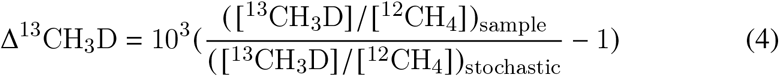

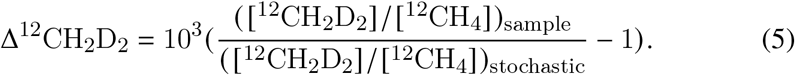

Here the bracketed values represent the concentrations of methane isotopologues. Equilibrium values of Δ^13^CH_3_D and Δ^12^CH_2_D_2_ used in this study are derived from the equilibrium constants in Young et al. (2017).

#### 2.3.2. Calculations of apparent fractionation and clumped isotopologue factors

Bulk isotope fractionation in a closed system is expected to follow the Rayleigh equation where the reactants (in this case CH_4_ gas) are well mixed throughout the process. Therefore calculating net fractionation factors follows the specific forms used here for methane ^13^C/^12^C and D/H after Mariotti et al. (1981):

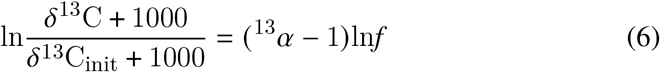

and

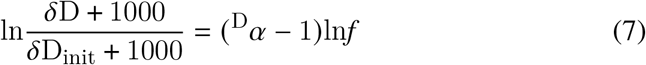

where the ^13^*α* and ^D^*α* are the net fractionation factors for ^13^C /^12^C and D /H, respectively. The *δ*^13^C and *δ*D are the bulk isotope values of residual methane (in ‰) at the time of sampling, and the *δ*^13^C_init_ and *δ*D_init_ are the values of methane (in ‰) at the start of the experiment. Here *f* is taken to be the fraction of methane remaining relative to the initial headspace methane. The net fractionation factors for the doubly-substituted isotopologues can be expressed similarly as:

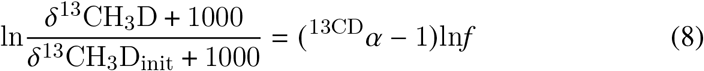

and

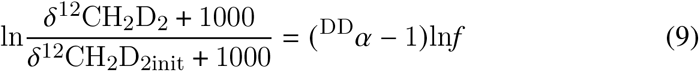

where ^13CD^*α* and ^DD^*α* are the net fractionation factors for ^13^CH_3_D and ^12^CH_2_D_2_, respectively, following prior studies (Wang et al., 2016; Liu et al., 2023).

These equations predict that if the isotope fractionations are Rayleigh-like, the relationship between observed fractionations (left hand sides of Equations 6, 7, 8, and 9) and corresponding ln*f* values are linear with with slopes of (*α* − 1). The net fractionation factors for our experiments, *α*, are therefore calculated from slopes defined by plots of observed fractionations and ln*f* values as determined by least-squares (York et al., 2004). The derived fractionation factors and 1*σ* uncertainties for the experiments at 37 °C, 27 °C and 21 °C are shown in Table 2. An important feature of both the 37 °C and 27 °C experiments is the change in net fractionation factors occuring at low *f* values (Figure 3), where the linearity of the regression is noticeably degraded if all data points are fit with a single line (Figure 3A, 3B, 3E, 3F). To address these changes, two separate sets of fractionation factors are calculated for the 37 °C and 27 °C experiments, respectively. The first set is calculated for the first five points in the 37 °C experiment, with *f* between 0.668 and 1.000. The second set is calculated for the last four points in the same experiment, with *f* ranging from 0.227 to 0.556 (Figure 3). For the 27 °C experiment, the first set of fractionation factors is calculated using the first four points with *f* values 0.374 and 1.000. Then the fourth data point (*f* = 0.374) and the last two points with lower *f* values are used for calculating the second set of fractionation factors (Figure 3A, 3B, 3E and 3F).

**Table 2:** Experimentally-derived net fractionation factors in this study and published data in Wang et al. (2016) and Krause et al. (2022). The numbers in parentheses represent the range of *f* for the data used in the calculation.

| Experiment | $^{13}\alpha$ | $1\sigma$ | $D\alpha$ | $1\sigma$ | $^{13}\text{CD}\alpha$ | $1\sigma$ | $^{13}\text{CD}\gamma$ | $1\sigma$ | $\text{DD}\alpha$ | $1\sigma$ | $\text{DD}\gamma$ | $1\sigma$ |
| --- | --- | --- | --- | --- | --- | --- | --- | --- | --- | --- | --- | --- |
| 37 °C ($f \geq 0.668$) | 0.9671 | 0.0011 | 0.6967 | 0.0101 | 0.6716 | 0.0110 | 0.9968 | 0.0218 | 0.4309 | 0.0193 | 0.8877 | 0.0474 |
| 37 °C ($f \leq 0.556$) | 0.9887 | 0.0003 | 0.8892 | 0.0028 | 0.8775 | 0.0031 | 0.9981 | 0.0047 | 0.7542 | 0.0064 | 0.9539 | 0.0101 |
| 27 °C ($f \geq 0.374$) | 0.9713 | 0.0024 | 0.7452 | 0.0209 | 0.7249 | 0.0227 | 1.0015 | 0.0422 | 0.5291 | 0.0387 | 0.9528 | 0.0878 |
| 27 °C ($f \leq 0.374$) | 0.9649 | 0.0049 | 0.6683 | 0.0466 | 0.6396 | 0.0507 | 0.9919 | 0.1048 | 0.3392 | 0.0929 | 0.7595 | 0.2334 |
| 21 °C | 0.9757 | 0.0002 | 0.7742 | 0.0022 | 0.7580 | 0.0024 | 1.0035 | 0.0043 | 0.5841 | 0.0043 | 0.9745 | 0.0091 |
| No-Cu ($f \leq 0.574$) | 0.9911 | 0.0005 | 0.8304 | 0.0093 | 0.8234 | 0.0097 | 1.0005 | 0.0163 | 0.6526 | 0.0191 | 0.9464 | 0.0349 |
| High-Cu ($f \geq 0.743$) | 0.9670 | 0.0018 | 0.7012 | 0.0161 | 0.6772 | 0.0175 | 0.9987 | 0.0346 | 0.4459 | 0.0303 | 0.9069 | 0.0744 |
| Krause et al. (20 °C) | 0.9848 | 0.0001 | 0.7265 | 0.0010 | 0.7141 | 0.0011 | 0.9981 | 0.0017 | 0.4757 | 0.0023 | 0.9013 | 0.0045 |
| Wang et al. (30 °C) | 0.9880 | 0.0003 | 0.8950 | 0.0030 | 0.8847 | 0.0030 | 1.0005 | 0.0003 | N/A | N/A | N/A | N/A |
| Wang et al. (37 °C) | 0.9780 | 0.0010 | 0.7980 | 0.0100 | 0.7804 | 0.0098 | 1.0000 | 0.0007 | N/A | N/A | N/A | N/A |

We also considered clumped isotopologue factors (*γ*), which Ono et al. (2021) utilize to describe deviations from the rule of the geometric mean:

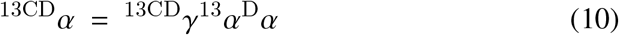

and

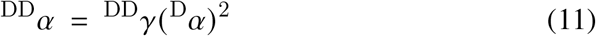

where *γ* values equal to unity means the frationation factors for the clumped isotopologues are simply the product of the fractionation factors for the heavy isotopes they contain (e.g. ^13CD^*α* = ^13^*α* ^D^*α*), consistent with the rule of the geometric mean (Bigeleisen, 1955). In this case, there is no energy difference in the formation of doubly substituted isotopologues compared with simple isotope substitution.

## 3. Results

Microbial growth rates presented across all experimental treatments in Figure 1A are calculated from OD_600_ measurements over time as shown in Figure B.9. Doubling times vary over an order of magnitude, from 7 to 113 hours. The longest doubling time (slowest growth rate) corresponds to the lowest growth temperature of 21 °C while the shortest doubling time (fastest growth rate) is obtained from the 37 °C experiments, consistent with the optimal conditions for this strain (Welander and Summons, 2012). Maximum OD_600_ values from temperature experiments range from ∼ 0.32 at 21 °C to ∼ 0.67 at 37 °C, respectively (Figure B.9). The growth profiles of the D-spiked experiments are similar to the unspiked 37 °C experiments.

**Figure 1:**
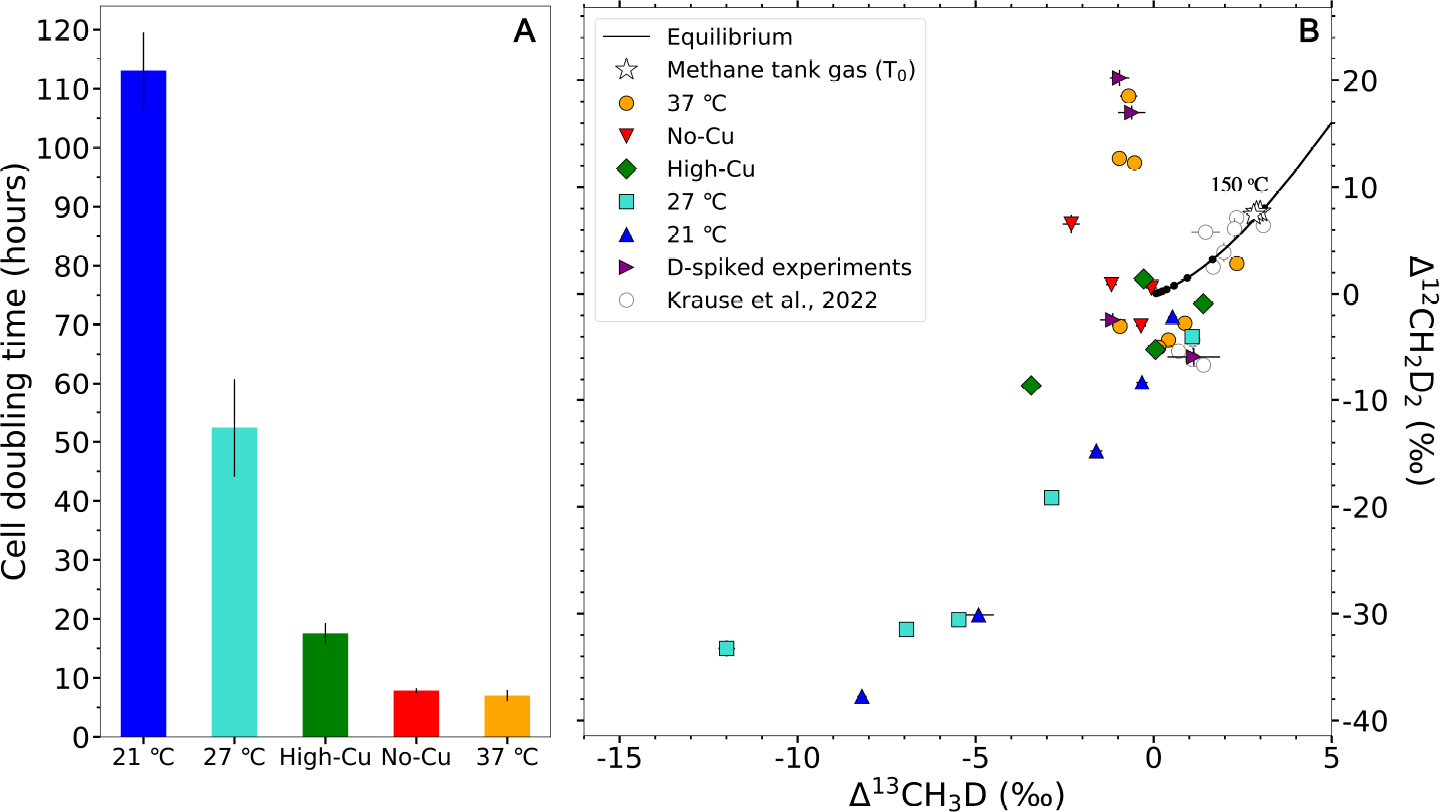
Cell doubling time of experimental culture (panel A) and clumped isotope signatures of headspace methane in Δ^12^CH_2_D_2_ vs. Δ^13^CH_3_D space (panel B). In panel A, longer cell doubling time represents slower growth rate. In panel B, the growth conditions are shown as growth temperature or total copper concentration in the legend. Experiments under different growth temperature are conducted with 5 *μ*M copper-supplemented media. Experiments with different copper concentrations are under 37 °C. High- and no-Cu correspond to 50 and 0 *μ*M Cu, respectively. The equilibrium curve in panel B is based on the equilibrium clumped isotope values in Young et al. (2017). The temperature increment between two adjacent points is 150 °C. The clumped isotope data from Krause et al. (2022) are shown for comparison as hollow circles.

Total headspace methane quantities are plotted in Figure B.9. The average amounts of methane consumed differed across experimental conditions. In 37 °C, 27 °C and no-Cu experiments, the average methane amount decreases from around 600 *μ*mol to less than 100 *μ*mol. On the other hand, high-Cu and 21 °C experiments have less total methane consumption, from around 600 *μ*mol to around 200 *μ*mol. The amount of methane in each serum bottle is also expressed as the fraction of residual methane (*f*) relative to the initial headspace methane, as shown in Table 1 and Figure 2. Dissolved methane only composes 1 ‰ of headspace methane in each sample, so it is neglected in the calculation of total methane. The lowest *f* value is 0.134, corresponding to 85.1 *μ*mol of headspace methane, which is approaching the minimum methane amount required for purification and mass spectrometry. Total methane oxidation rate increases from 8.89 to 30.88 *μ*mol/hr and 1.94 to 3.79 *μ*mol/hr in 37 °C and 27 °C experiments, respectively (Figure B.9). In contrast, oxidation rate in the 21 °C experiment stays constant at 0.96 *μ*mol/hr (Figure B.9).

**Figure 2:**
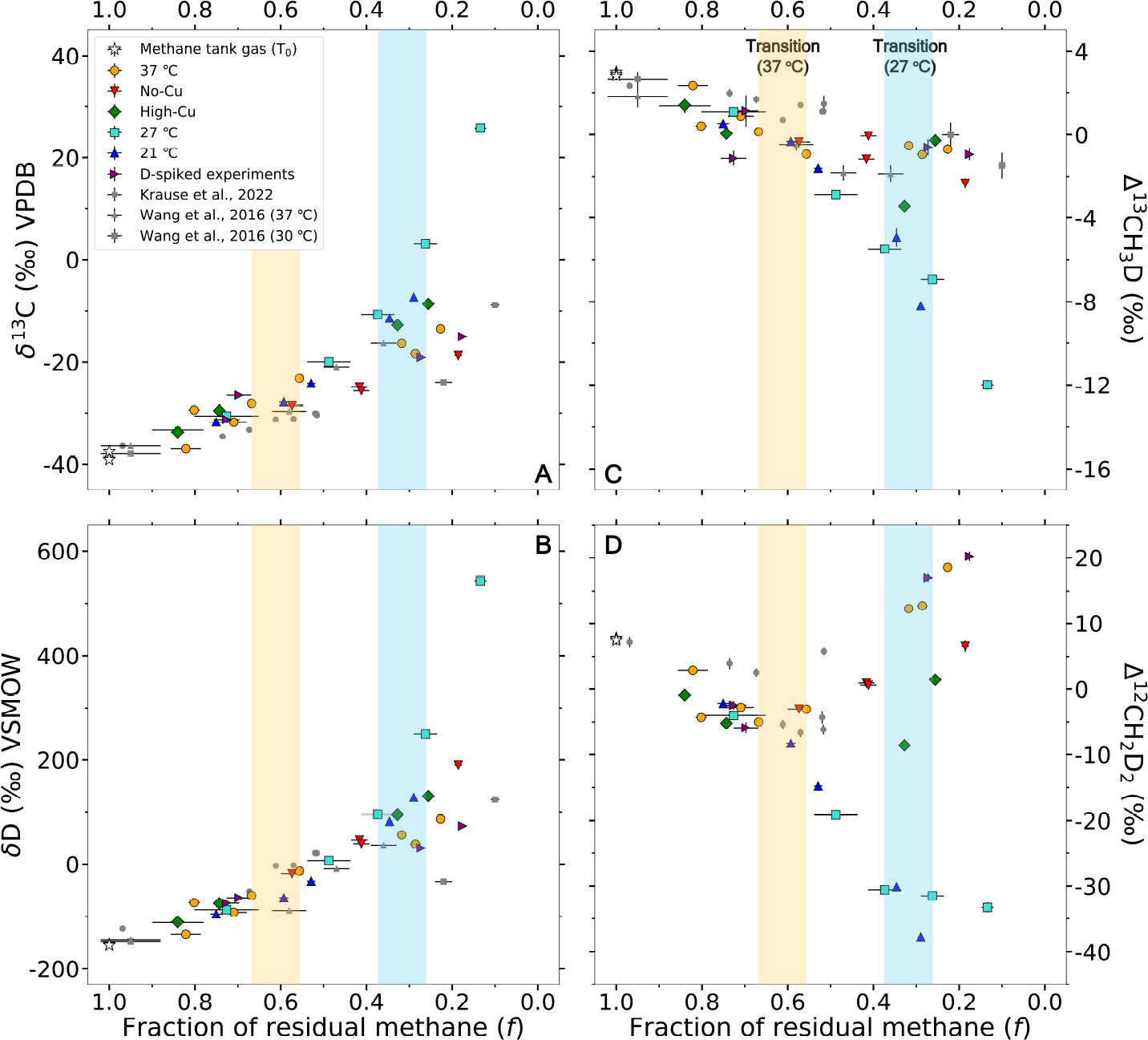
Variations of isotopic compositions with the fraction of headspace methane remaining after oxidation (*f*) : A) *δ*^13^C ; B) *δ*D ; C) Δ^13^CH_3_D ; D) Δ^12^CH_2_D_2_. In each panel, lower *f* values (the right side) represent more methane consumption. The light yellow and the blue bars show the range of *f* values where the isotope signatures start to deviate from the Rayleigh fractionation in 37 °C and 27 °C experiments. Data from previous studies (Wang et al., 2016; Krause et al., 2022) are included as the gray data points in the figure.

Dissolved copper concentration ([Cu^2+^] _aq_) in the media range from 0.043 ± 0.026 to 0.864 ± 0.029 *μ*M in 5 *μ*M Cu-supplemented experiment, with lower [Cu^2+^] _aq_ towards the end of the experiment. The [Cu^2+^] _aq_ ranges from 7.54 ± 0.06 to 9.19 ± 0.06 *μ*M in 50 *μ*M (high) Cu-supplemented experiment, with no significant decrease towards the end of the experiment. The [Cu^2+^] _aq_ of the no-Cu experiment is around zero throughout the experiment, considering 1*σ* error (Figure B.10).

The measured *δ*^13^C of all samples are between −39.10 ± 0.01 to 25.84 ± 0.02 ‰ (1*σ*, same thereafter) VPDB and *δ*D are in the range of −153.97 ± 0.02 to 543.32 ± 0.09 ‰ VSMOW (Table 1, Figure 2A, 2B). Consistent enrichment of heavier isotopes (^13^C and D) in residual methane is observed in all experiments. The average Δ^13^CH_3_D and Δ^12^CH_2_D_2_ values for the methane tank gas are 2.90 ± 0.09 ‰ and 7.6 ± 0.2 ‰ (Table 1, Figure 1B), which are close to the equilibrium values at 165 °C. The measured Δ^13^CH_3_D values from all experiments range from −11.99 ± 0.23 ‰ to 2.99 ± 0.19 ‰ while Δ^12^CH_2_D_2_ range from −37.7 ± 0.6 to 20.2 ± 0.8 ‰ (Table 1, Figure 1B). Δ^13^CH_3_D values consistently decrease with more methane consumption in all experiments, except for the 37 °C and high-Cu experiments (Figure 2C). More prominent decrease of Δ^13^CH_3_D values are observed at lower growth temperatures. Similarly, Δ^12^CH_2_D_2_ values decrease with methane consumption at lower growth temperatures (21 and 27 °C). In contrast, in the experiments at 37 °C, Δ^12^CH_2_D_2_ values decrease slightly at high methane concentrations (*f* > 0.5) and then increase at low methane concentrations (*f* < 0.5), as shown in Figure 2C and 2D. The measured *δ*D for the media of the D-spiked experiments ranges from 1419 ± 3 to 1440 ± 3 ‰, and from 3003 ± 6 to 3017 ± 7 ‰ in two sets of D-spiked experiments, respectively (Table C.5). The average values across biological replicates are 1433 ± 10 ‰ (1 standard deviation) and 3008 ± 6 ‰. The isotope signatures of the two D-spiked experiments follow the same pattern as the non-spiked experiment at 37 °C, so they are discussed together as one set of experiment in the following sections.

Net fractionation factors across all experiments range from 0.9649 ± 0.0049 to 0.9911 ± 0.0005 for ^13^*α* (Figure 3A, 3C), from 0.6683 0.0466 to 0.8892 ± 0.0028 for ^D^*α* (Figure 3B, 3D), from 0.6396 ± 0.0507 to 0.8775 ± 0.0031 for ^13CD^*α* (Figure 3E, 3G), and from 0.3392 ± 0.0929 to 0.7542 ± 0.0064 for ^DD^*α* (Figure 3F, 3H). The net fractionation factors shift towards unity at low methane concentration for the 37 °C experiments (Table 2; Figure 3) regardless of copper concentrations, indicating less discrimination between heavy to light isotopologues at low *f* values, as methane was depleted. In contrast, the 27 °C experiment exhibits more net fractionation at low methane concentration (Table 2; Figure 3A, 3B, 3E, 3F), such that the reaction shows preference for the light isotopes or isotopologues late into the reaction coordinate. The 21 °C experiment shows a consistent net fractionation over the whole experiment (Table 2; Figure 3A, 3B, 3E, 3F). The net fractionation during the early stages of all experiments are comparable (Table 2), as reflected by the similarities in the trends of isotope signatures in the 37 °C (*f* ≥ 0.668), 27 °C (*f* ≥ 0.374), 21 °C and high-Cu (*f* ≥ 0.743) experiments (Figure 1B; Figure 2). During the early stages of all experiments, the net fractionation factors scale inversely with temperatures.

**Figure 3:**
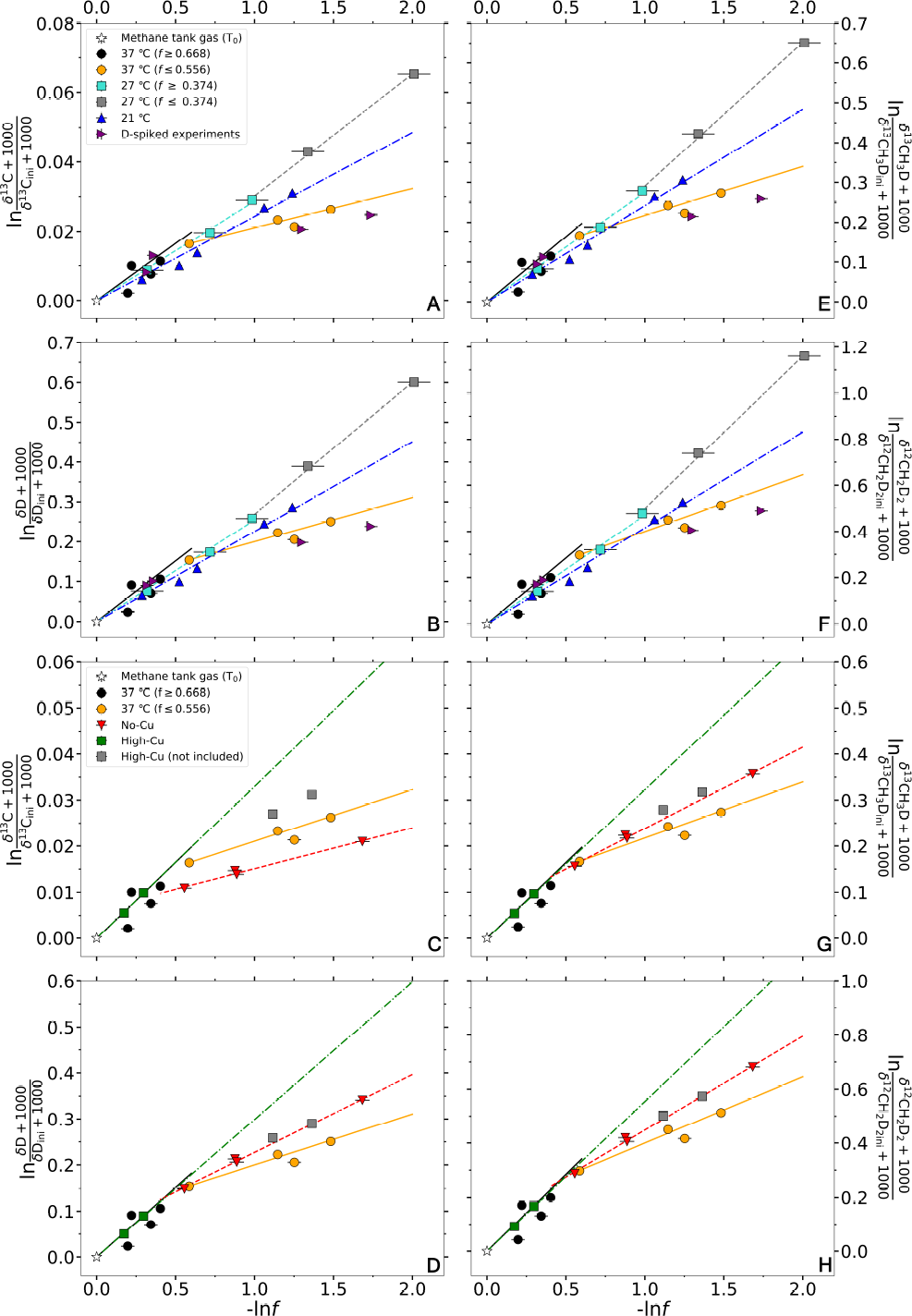
Linear regressions for calculating net fractionation factors for different temperatures, water spikes and copper concentrations. The slopes of the regression lines are 1-*α*, where *α* is the net fractionation factor of the isotope or isotopologue assuming a Rayleigh fractionation process (values are shown in Table 2). Panels A to D show the ^13^*α* and ^D^*α* across temperatures and copper concentrations; panels E and H show the ^13CD^*α* and ^DD^*α* for different temperatures and copper concentrations. The right side of each panel represents lower *f* values. Within the 37 °C experiment (orange and black circles in panels A, B, E, F), the net fractionation of *f*≤ 0.556 (orange circles) is significantly different from *f* ≥ 0.668 (black circles). In the 27 °C experiment (gray and cyan squares in panels A, B, E, F), the net fractionation is also different between *f* ≤0.374 (gray squares) and *f* ≥ 0.374 (cyan squares). The isotope signatures of deuterium-spiked experiments (purple triangles in panels A, B, E, F) are not significantly different from non-spiked experiments at the same temperature, regardless of the spike. Two data points (gray rectangles in panels C, D, G, H) in the high-Cu experiment are not included in the regression, as they deviate from the Rayleigh curve determined by the first three points, and they are not enough to do a separate linear regression. Regardless of copper concentrations, deviation from Rayleigh fractionation still occurs at low *f* (panels C, D, G, H).

## 4. Discussion

### 4.1. Net isotope fractionation factors

The bulk isotope fractionation factors ^13^*α* and ^D^*α* broadly agree with the direction and magnitude of those previously reported (Feisthauer et al., 2011; Rasigraf et al., 2012). This said, we observe more D/H fractionation, with the 37 °C (*f* ≥ 0.668) and 27 °C (*f* ≤0.374) experiments yielding ^D^*α* = 0.6967 ± 0.0101 and 0.6683 ± 0.0466, respectively, whereas the previously estimated D/H fractionation values for MMO range from 0.7685 ± 0.0303 to 0.8900 ± 0.0115 (Feisthauer et al., 2011; Rasigraf et al., 2012). The range of isotope fractionation factors in the 37 °C experiment encompass the factors from Wang et al. (2016) at the same temperature (Table 2). The fractionation factors in our 21 °C experiment with *M. capsulatus* agree with the values obtained at room temperature (∼ 20 °C) using a different strain, *Methylosinus trichosporium* OB3b (Krause et al., 2022). This is reasonable since both strains possess pMMO and sMMO for methane oxidation. Despite the deviation from purely Rayleigh fractionation, ^13CD^*γ* in all our experiments are close to unity within 1*σ* error (Table 2), consistent with previous reports (Wang et al., 2016; Krause et al., 2022). On the other hand, ^DD^*γ* shows large differences across the experimental conditions tested, ranging from 0.7595 ± 0.2334 to 0.9745 ± 0.0091. The highest ^DD^*γ* value (0.9745 ± 0.0091), occurring in the 21 °C experiment, is higher than the value from Krause et al. (2022) (0.9013 ± 0.0045) at a similar growth temperature, indicating less deviation from the rule of geometric mean. In contrast, in the 37 °C (*f* ≥ 0.668) experiment, the ^DD^*γ* is 0.8877 ± 0.0474, which is closer to the value from Krause et al. (2022), albeit at different temperatures. The 27 °C (*f*≤ 0.374) experiment has the lowest ^DD^*γ* value (0.7595 ± 0.2334), showing the most deviation from the rule of geometric mean.

One prominent feature in our experiments is the change of net isotope fractionation during the progressive oxidation of methane (Figure 3, Table 2). Similar changes in bulk carbon isotope fractionation are reported by Templeton et al. (2006), and attributed to an increase in the total methane oxidation rate relative to the rate of methane conversion from the gaseous to the dissolved state and diffusion into the cells. In another study, the variation of net carbon isotope fractionation with temperature was attributed to the ratio of adsorption and desorption of methane onto the cell wall and the rate of conversion from methane to methanol (Nihous, 2010). To explore this possibility in our experiments, we investigate three scenarios that could generate the observed changes in apparent fractionation factors within the three differential temperature experiments. These scenarios are i) differential enzyme isotope effects as a function of dissolved copper, ii) isotopic re-equilibration between methane and water, and iii) reservoir effects related to the ratio between the rate of methane transport across the cell membrane relative to the total methane oxidation rate.

#### 4.1.1. Enzymatic effect on net fractionation

It is possible that the changes in the net fractionation factor originate from the switch of methane oxidation pathways. The first step in AeOM (from methane to methanol) is catalyzed by two isoforms of the methane monooxygenase enzyme, the particulate methane monooxygenase (pMMO) and the soluble methane monooxygenase (sMMO) (Sirajuddin and Rosenzweig, 2015; Ross and Rosenzweig, 2017). Key differences between the two enzymes include either a monocopper, dicopper, or zinc/copper active site in pMMO, whereas sMMO possesses only a diiron active site (Sirajuddin and Rosenzweig, 2015). The oxygen activation mechanism by the diiron center of sMMO is well studied (Gassner and Lippard, 1999), while the catalytic mechanism behind pMMO is less well characterized (Koo and Rosenzweig, 2021). The *M. capsulatus* genome encodes both pMMO and sMMO (Ribbons and Michalover, 1970; Colby et al., 1977), and the expression of pMMO versus sMMO proteins is controlled by environmental concentrations of copper. High copper concentrations induce the organism to express pMMO and therefore lead to high pMMO activity, whereas limited or no copper induces cells to express sMMO – this is known as the the “copper switch” (Stanley et al., 1983; Prior and Dalton, 1985; Nielsen et al., 1997; Koo and Rosenzweig, 2021).

In this study, progressive and substantial cellular uptake of dissolved copper occurs during population growth, which is indicated by the significant drop of [Cu^2+^]_aq_ in the 37 °C experiments with 5 *μ*M initial dissolved copper (Figure B.10). It is worth noting that the maximum [Cu^2+^]_aq_ in both Cu-supplemented experiments are lower than the total Cu added by five to six fold due to the precipitation of copper salts. However, the loss of [Cu^2+^] _aq_ due to microbial uptake is replenished by the re-dissolution of Cu-bearing precipitates, which is supported by the observation that the [Cu^2+^] _aq_ remains above 7 *μ*M over the entirety of the high-Cu experiment (Figure B.10). The [Cu^2+^] _aq_ in the media changes from abundant at the beginning to depleted at the end of the 37 °C experiments with 5 *μ*M Cu due to microbial uptake, which could trigger the “copper switch”. In contrast, cells in the no-Cu experiment are always Cu-limited and therefore expressing sMMO, and that the high-Cu experiment is never Cu-limited and so cells express primarily pMMO.

In principle, it is possible that the change in net fractionation factors is a result of the change in the activity of pMMO versus sMMO. However, the change in net fractionation occurs in both the no-Cu and high-Cu experiments (Figure 3C, 3D, 3G, 3H), indicating that the shift in net fractionation is unlikely to result from enzyme-specific effects. In the high-Cu experiment, the calculated net fractionation factors using data points with *f*≥ 0.743 are similar to the result of the 37°*C* (*f* ≥ 0.668) experiment (Table 2). The two data points with *f* ≤ 0.328 show the same isotopic depletion relative to the regression line derived from the points with higher *f* values, which is the same as the trend in the 37 °C experiment (Figure 3C, 3D, 3G, 3H), so are not included in the regression. In the no-Cu experiment, even though the highest *f* value for the data points is 0.574 (Table 1, Figure 2), which is the lower boundary of *f* values where fractionation factors change, the corresponding net fractionation factors calculated without time zero are very similar to 37 °C (*f* ≤0.556) (Table 2, Figure 3C, 3D, 3G, 3H).

The change in net fractionation factors correlates with the quantity of residual methane, not the change in copper availability. We show that the clumped isotope fractionation values are similar for pMMO-dominant (high-Cu) and sMMO-dominant (no-Cu) conditions, and that pMMO and sMMO do not have unique bulk carbon and hydrogen isotope fractionation factors, consistent with prior work (Feisthauer et al., 2011). It is therefore unlikely that enzyme-specific effects are significant contributors to the isotopic trajectories we observe for AeOM.

#### 4.1.2. Hydrogen isotope exchange with water

Another putative mechanism for the observed divergence from Rayleigh fractionation (Figure 3) is hydrogen isotope exchange between methane and water. In contrast to AOM, the first step of AeOM is generally considered irreversible and dominated by large kinetic isotope fractionations (Lee et al., 1993; Nesheim and Lipscomb, 1996). If isotope exchange between water and methane were to occur, the *δ*D of methane should move towards the equilibrium values with water during the experiment. At 37 °C, the equilibrium fractionation factor between water and methane 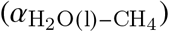 is 1.260 (Horibe and Craig, 1995), which translates to 931 and 2181 ‰ for methane equilibrated with water having *δ*D of 1433 and 3008 ‰, respectively. The observed *δ*D of methane in the D-spiked experiments are well below the equilibrium values (Table 2). The other isotope signatures (*δ*^13^C, Δ^13^CH_3_D and Δ^12^CH_2_D_2_) of the D-spiked experiments are all similar to the results from the non-spiked 37 °C experiments (Figure 2A, 2C, 2D). Therefore, we rule out the possibility of hydrogen isotope exchange with water.

#### 4.1.3. Reservoir effect linked to methane transport and consumption

With the enzymatic effect and methane-water isotopic exchange ruled out, we consider a third potential mechanism - the reservoir effect in a closed-system. In other words, we consider that headspace methane and intracellular methane are distinct reservoirs. The measured isotopic signals are those for the headspace methane, the first reservoir, while the observed fractionation factors result from the combination of cross-membrane methane transport and methane oxidation, involving transport between the reservoirs and loss of methane to the second reservoir, respectively.

In order to examine this possibility, we construct and solve a two-box model to simulate the cross-membrane transport between methane inside and outside of the cells (Figure 4). Following Nihous (2010), we assume that the cross-membrane transport (*J*_in_ and *J*_out_) does not fractionate isotopes, but we do consider the possibility that rates change over the duration of the experiment. Methane is oxidized inside the cell, with an oxidation rate of *J*_oxidize_. We assume that the intra-cellular oxidation has associated net isotope fractionation factors corresponding to those calculated at high *f* values, where reservoir effects are likely minimal. For simplicity, we assume methane concentrations inside the cells are at steady state, such that:

**Figure 4:**
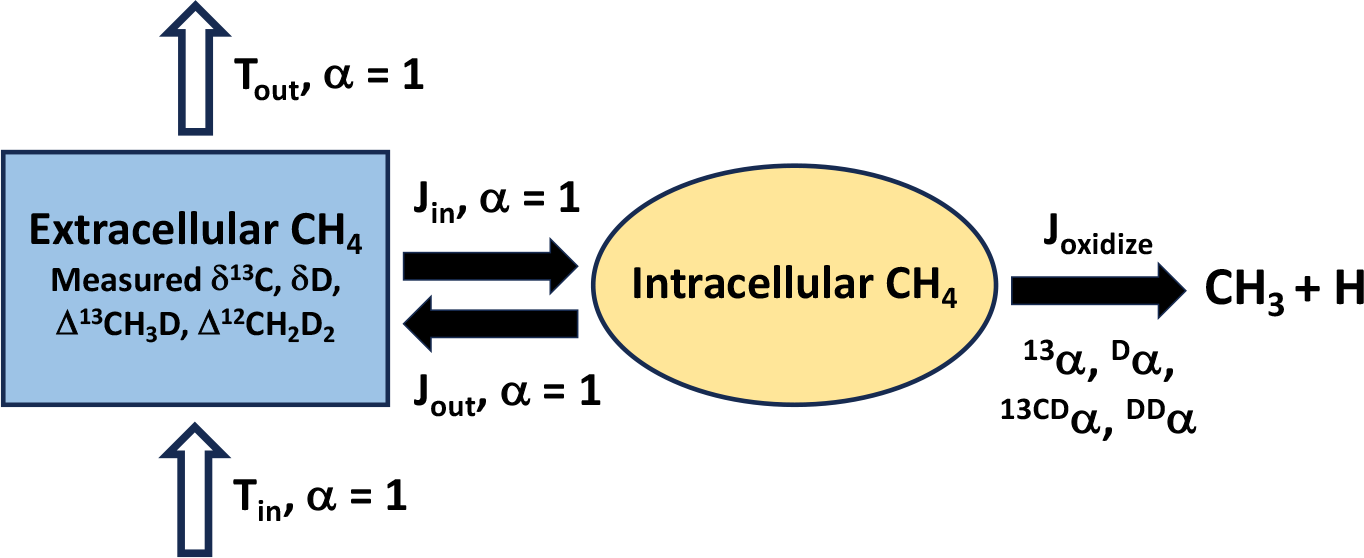
An illustration of the two-box model developed in this study. The closed-system model only includes the solid arrows while the open-system model includes both the solid and hollow arrows. Methane is distinguished as extracellular methane (blue box) and intracellular methane (yellow box). The measured isotope signatures are the isotope compositions of extracellular methane. Cross-membrane transport rates are marked as *J*_in_ and *J*_out_ and assumed to have no isotope fractionation. Methane oxidation rate is represented by *J*_oxidize_ and have the isotope fractionation derived from the experiments. The open-system model has additional advection input and output rates (*T*_in_ and *T*_out_, shown as the hollow arrows) without isotope fractionation.

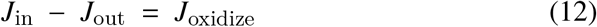

For AeOM, only the first step of oxidation is considered since the reversibility of this step is low, suggesting that it is responsible for the rate of oxidation and associated kinetic isotope effects (Nesheim and Lipscomb, 1996). We adopt a reaction network similar to that utilized for AOM in Liu et al. (2023) (Table 3), with the modification that reactions are all assumed to be irreversible. This assumption is supported by the D-spiked experiments, as well as previous studies of AeOM (Lee et al., 1993). Therefore, only the reactions that break C-H or C-D bonds are accounted for in Table 3. The experimentally-derived net fractionation factors represent the ratios of rate constants between the heavy isotopologues and ^12^CH_4_. The measured net D/H fractionations are weighted convolutions of both primary and secondary fractionation effects (Ono et al., 2021; Liu et al., 2023). Due to the lack of data distinguishing primary and secondary isotope effects, we do not make this distinction in our reaction network. The primary and secondary fractionations are therefore taken to be the same in our network (e.g. Table 3, reactions 2 and 3) in order to match the experimental results. This use of a single, effective *α* is equivalent to using weighted sums of distinct primary and secondary fractionation factors and has no effect on the final outcome, as discussed by Liu et al. (2023). The net fractionation factors from 37 °C (*f* ≥ 0.668), 27 °C (*f* ≥ 0.374) and 21 °C are taken as representative of the fractionation factors absent reservoir effects, and they are assumed to be constant throughout each experiment. A set of ordinary differential equations (ODEs) is constructed to describe the time variation of methane isotopologues inside and outside the cells, and the numerical solutions are obtained at each time step (detailed in the Appendix). The temporal change of abundances are then translated into the changes in isotopologue ratios with *f*.

**Table 3:** Reactions, rate constants and fractionation factors used to calculate isotope fractionations during AeOM. The set of fractionation factors used for the 37 °C experiment is given as an example in the table. *K*_oxidize_ is a tunable coefficient for controlling the oxidation rate *J*_oxidize_. The relation between *K*_oxidize_ and *J*_oxidize_ is in the Appendix.

| Reaction | Rate constant | Fractionation factor |
| --- | --- | --- |
| $^{12}\text{CH}_4 \rightarrow ^{12}\text{CH}_3 + \text{H}$ | $k_1 = K_{\text{oxidize}} \alpha_1$ | $\alpha_1 = 1.0000$ (1) |
| $^{12}\text{CH}_3\text{D} \rightarrow ^{12}\text{CH}_3 + \text{D}$ | $k_2 = 1/4 K_{\text{oxidize}} {}^{\text{D}}\alpha$ | ${}^{\text{D}}\alpha = 0.6967$ (2) |
| $^{12}\text{CH}_3\text{D} \rightarrow ^{12}\text{CH}_2\text{D} + \text{H}$ | $k_3 = 3/4 K_{\text{oxidize}} {}^{\text{D}}\alpha$ | ${}^{\text{D}}\alpha = 0.6967$ (3) |
| $^{12}\text{CH}_2\text{D}_2 \rightarrow ^{12}\text{CH}_2\text{D} + \text{D}$ | $k_4 = 1/2 K_{\text{oxidize}} {}^{\text{DD}}\alpha$ | ${}^{\text{DD}}\alpha = 0.4309$ (4) |
| $^{12}\text{CH}_2\text{D}_2 \rightarrow ^{12}\text{CHD}_2 + \text{H}$ | $k_5 = 1/2 K_{\text{oxidize}} {}^{\text{DD}}\alpha$ | ${}^{\text{DD}}\alpha = 0.4309$ (5) |
| $^{13}\text{CH}_4 \rightarrow ^{13}\text{CH}_3 + \text{H}$ | $k_6 = K_{\text{oxidize}} {}^{13}\alpha$ | ${}^{13}\alpha = 0.9671$ (6) |
| $^{13}\text{CH}_3\text{D} \rightarrow ^{13}\text{CH}_3 + \text{D}$ | $k_7 = 1/4 K_{\text{oxidize}} {}^{13\text{CD}}\alpha$ | ${}^{13\text{CD}}\alpha = 0.6716$ (7) |
| $^{13}\text{CH}_3\text{D} \rightarrow ^{13}\text{CH}_2\text{D} + \text{H}$ | $k_8 = 3/4 K_{\text{oxidize}} {}^{13\text{CD}}\alpha$ | ${}^{13\text{CD}}\alpha = 0.6716$ (8) |

We conducted a sensitivity test to examine how the ratio of the transport rate *J*_in_ relative to the oxidation rate *J*_oxidize_ can influence the trajectory of isotope signatures. We change the value of either *J*_in_ or *J*_oxidize_ while keeping the other constant (Figure 5). To match the scale of experimental data, we use units of *μ*mol/hr for our rates. In the first test, we use *J*_in_ values of 500, 50, 30, 20, 10 and 5 *μ*mol/hr at a fixed *J*_oxidize_ of 3.25 *μ*mol/hr (corresponding to *K*_oxidize_ = 1 in Table 3). In the second test, *J*_in_ is fixed at 20 *μ*mol/hr, while *J*_oxidize_ varies, with values of 0.13, 1.30, 2.16, 3.25, 6.50 and 13.0 *μ*mol/hr. Therefore, the same set of *J*_oxidize_/*J*_in_ ratios are used in the two tests. This ratio can be described as a Damköhler number (Da) (Bailey et al., 2018) such that

**Figure 5:**
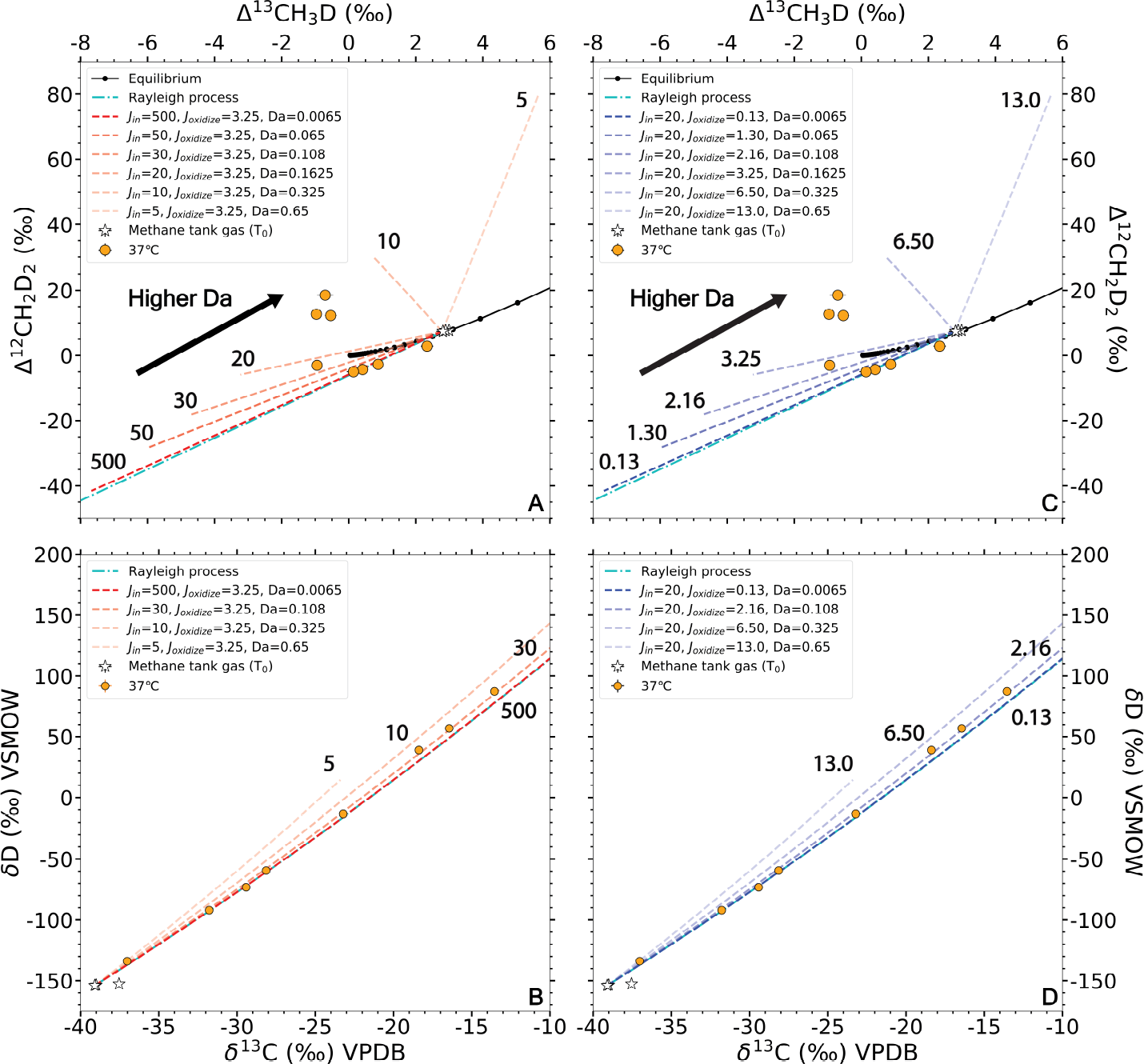
Oxidation trajectories with different but constant *J*_in_ and *J*_oxidize_. Panels A and B are oxidation trajectories with varying *J*_in_ and constant *J*_oxidize_ in clumped and bulk isotope phase spaces. Panels C and D are oxidation trajectories with varying *J*_oxidize_ and constant *J*_in_ in clumped and bulk isotope phase spaces. Oxidation trajectories of pure Rayleigh processes are shown as cyan dashed lines. The numbers by the trajectories mark the values of varying rates. The arrows in panel A and C point to the direction of increasing Da. Variations between Da are less significant in bulk isotope phase space, thus only four curves with Da = 0.0065, 0.108, 0.325 and 0.65 are shown in panel B and D.

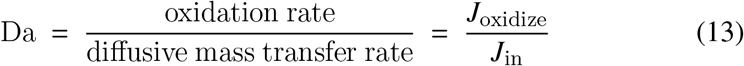

where Da is a dimensionless number between 0 and 1, and higher Da values represent more methane oxidation relative to cross-membrane transport of methane. The oxidation trajectories with Da = 0.0065, 0.065, 0.108, 0.1625, 0.325 and 0.65 are shown in Figure 5. For comparison, we use the equation described in Wang et al. (2016) and Liu et al. (2023) to calculate the oxidation trajectory of a closed-system Rayleigh process as a function of *f* :

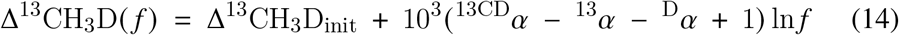

and

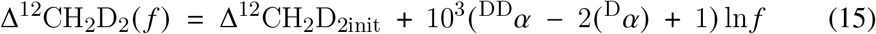

where Δ^13^CH_3_D_init_ and Δ^12^CH_2_D_2init_ are the initial clumped isotope signatures of methane, and Δ^13^CH_3_D (*f*) and Δ^12^CH_2_D_2_ (*f*) are the clumped isotope signatures of methane at a specified value for *f*.

The oxidation trajectories that share the same Da have the same pattern in both bulk and clumped isotope phase space (Figure 5), illustrating that the relative magnitude of methane oxidation rate versus transport rate is the determining factor of the oxidation trajectories in this model. However, it is worth noticing that at a low methane oxidation rate, the total time to reach the same *f* is longer, which makes the reaction time an important factor to account for when fitting the experimental data. When the transport dominates the system (low Da), the oxidation trajectory is closer to a purely Rayleigh process. This means the isotopically fractionated methane inside the cell is immediately expelled from the cell so that the isotopic composition of methane outside reflects only the isotope fractionation of the oxidation process. Conversely, in a transport-limited system (high Da), strongly fractionated methane gas is not immediately transported out of the cell, resulting in deviations from Rayleigh fractionation trajectories (i.e., there is a reservoir effect). As the transport rates decrease to values of the same order of magnitude as the oxidation rate (e.g. *J*_in_ decreases from 500 to 5 *μ*mol/hr, and Da number closer to 1), the trajectory deviates further from Rayleigh fractionation, eventually leading to a completely different direction in clumped isotope phase space (Figure 5A and 5C). Departures from Rayleigh distillation are more significant in clumped isotope phase space (Figure 5A and 5C) than in the bulk isotope phase space (Figure 5B and 5D). Similar changes in net isotope fractionation factors that originate from changes in relative rates of transport and consumption have been reported for bulk carbon isotope fractionation during microbial aerobic methane oxidation (Templeton et al., 2006) and for S isotopes during sulfate reduction (Wing and Halevy, 2014; Sim et al., 2017).

The results from the box model indicate that varying Da over the course of the experiments is required to fit the oxidation trajectories in the 37 °C and 27 °C experiments. To illustrate, the *J*_oxidize_ in eqn. 13 is set as the total methane oxidation rate calculated from the experimental data (Figure B.9). We then assume that the cross-membrane transport is a diffusion process in which the rate of transport follows Fick’s law. Given that *J*_in_ is not constrained by available experimental data, the Da is parameterized using the following equation adopted from Clark et al. (2014):

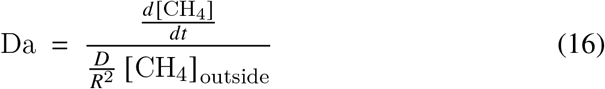

where *d* [CH_4_]/*dt* is the experimentally-derived total methane oxidation rate in *μ*mol/hr, and [CH_4_]_outside_ is the measured headspace methane concentration. The cross-membrane diffusion rate is parameterized by the ratio of methane diffusivity (*D*) to the square of the radius of the cells (*R*), as well as to the headspace methane concentration. The temporal change in Da is attributable to changes in the radii of the cells as well as in the methane concentration. Changes in *R* in this model, the collective radii of cells expressed as a single value, are constrained by the OD_600_ measurements. By assuming the volumes of the cells are 4/3*πR*^3^ and proportional to OD_600_, the effective radius *R*^2^ is proportional to OD^2/3^. In contrast, diffusivity (*D*) values are assumed to be constant. Since the ratio *D* /*R*^2^ (with the unit of s^−1^) is the most relevant for our model, rather than specific values for *R* and *D* where the length scales cancel, we convert the term *D*/ *R*^2^ to *C* /OD^2/3^, where *C* is a constant that is proportional to *D*. The values of *C* in the models are obtained from the best fit for the experimental data (Figure 6).

**Figure 6:**
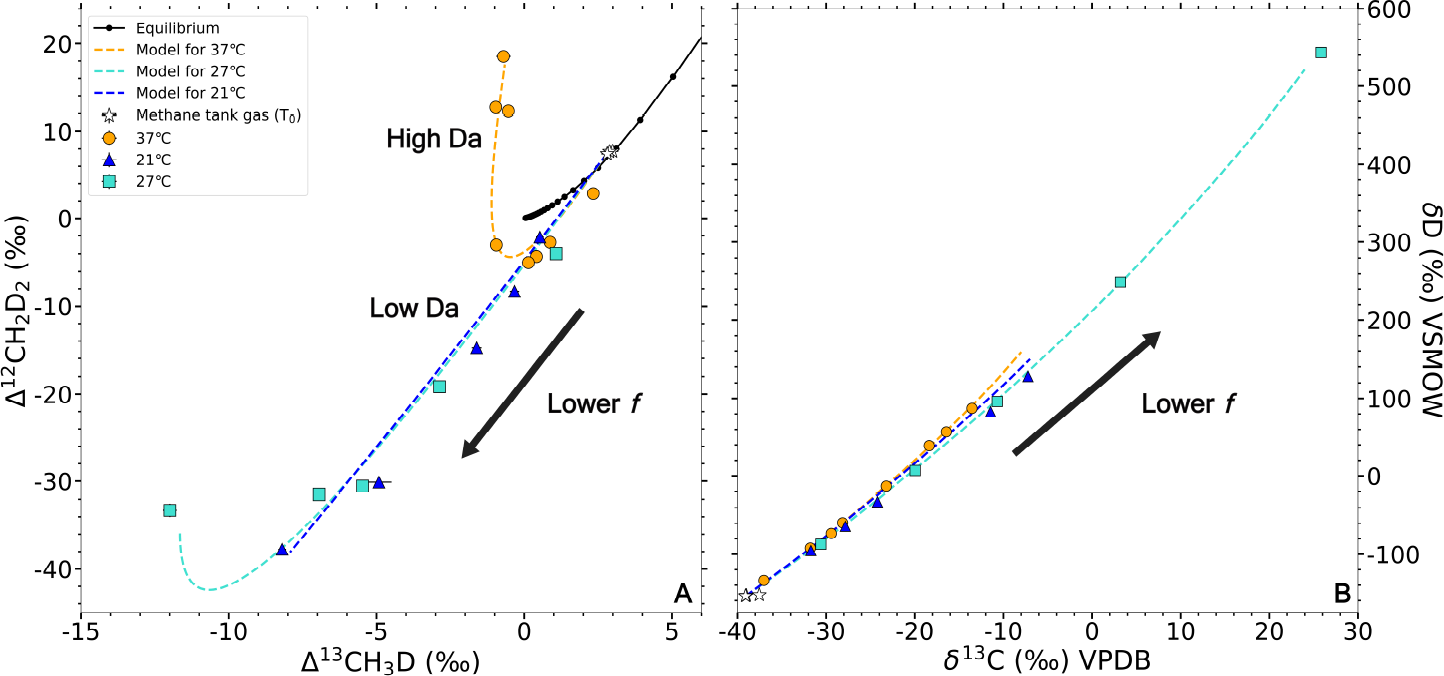
Modeled oxidation trajectories in clumped (panel A) and bulk (panel B) isotope phase space. Data from the 37 °C, 27 °C and 21 °C experiments are fitted by the model. The parameterized *J*_oxidize_, *J*_in_ and Da used for each curve are shown in Figure B.11. The increases in Δ^12^CH_2_D_2_ values with *f* are due to increases in Da.

The model curves in bulk and clumped isotope phase space are shown in Figure 6. The best-fit values for *C* are 0.19 and 0.08 for 37 °C and 27 °C experiments, respectively. The 21 °C experiment shows a Rayleigh-like oxidation trajectory, so any *C* value greater or equal to 0.04 fits the data (Figure 6). Despite this uncertainty, it is reasonable to conclude that the value of *C* decreases as temperature decreases. This indicates that *D* also decreases with temperature, consistent with the normal temperature dependence of diffusivity in general (Cussler, 2009). The model values for Da increase from 0.003 to 0.688, 0.0001 to 0.769, and 0.002 to 0.045 in 37 °C, 27 °C and 21 °C experiments, respectively (Figure B.11). This is due to a decrease in *J*_in_, as well as an increase in *J*_oxidize_ with microbial growth. *J*_oxidize_ in the model increases linearly to a maximum of 3.79 *μ*mol/hr at 27 °C, while it increases as a fourth-order polynomial with time to a maximum of 30 *μ*mol/hr at 37 °C (see Appendix). These changes in rate are used to make the model integrated average methane oxidation rate match the observed total methane oxidation rate. In the 21 °C model, *J*_oxidize_ is kept at a constant value of 0.96 *μ*mol/hr since the observed total methane oxidation rate is constant. Values for *J*_in_ decrease with time in all three models, as both the increase of OD (cell radii) and the decrease of headspace methane contribute to the decrease of transport rate.

In summary, the model indicates that in the 37 and 27 °C experiments, the magnitudes of oxidation rate are much smaller than transport rates at the beginning, but they become comparable to transport rates by the end of the experiments. In contrast, in the 21 °C experiment, the system remains dominated by transport rate throughout due to a low oxidation rate at the lower temperature.

### 4.2. Open system model with different cross-membrane transport

In natural environments, aerobic microbial methane oxidation often occurs in an open system. An example of such a system is methane diffusing up a stratified lake water column as described in Giunta et al. (2022). The Δ^13^CH_3_D and Δ^12^CH_2_D_2_ signatures of AeOM in an open-system with advection show different trends than those during closed-system AeOM. The influences of *γ* and the ratio between oxidation and advection (*ϕ*) were initially investigated by Wang et al. (2016) and Krause et al. (2022). Following the example of these previous works, we define *ϕ* as the ratio of methane oxidation rate and total advective input rate (Figure 4):

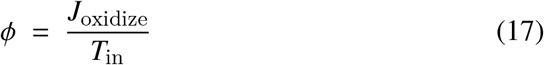

In this formulation, *ϕ* = 1 represents a fully oxidative system in which all methane gas advected into the system is oxidized, while *ϕ* = 0 represents a fully advective system in which all methane gas flows through the system without oxidation. The basic assumptions in the open-system model are: 1) the advection input (*T*_in_ in Figure 4) has the same isotopic composition as the initial headspace methane; and 2) there is no isotope fractionation during advection. Based on the ODE set described in *section 4*.*4*, we construct a new ODE set by adding two terms for advection in and out of the system, and the numerical solutions are obtained as before. Since input and output rates are fixed, steady states between the reaction and the flow are attained. For the methane oxidation, we use the fractionation factors derived from the 37 °C experiment. We test the effect of the Damköhler number discussed in *section 4*.*4* on the steady-state clumped isotope signatures at different *ϕ* values (Figure 7). The curves in Figure 7 are not individual time series, but rather each point on the curve represents a steady-state clumped isotope signature in an open-system scenario with specific values for Da and *ϕ*. At the same Da, the effect of *ϕ* on clumped isotope ratios closely resembles the effect described in Krause et al. (2022), with higher *ϕ* values resulting in more positive Δ^12^CH_2_D_2_ values at steady state. Conversely, the effect of higher Da values at a fixed *ϕ* value is a more positive Δ^13^CH_3_D value. On the other hand, the effect of Da on Δ^12^CH_2_D_2_ is not unidirectional, marked by a less range of variation in Δ^12^CH_2_D_2_ value at high Da. However, the effect of Da is minor compared with *ϕ* and the general trend of higher *ϕ* having more positive Δ^12^CH_2_D_2_ values is not affected by different Da.

**Figure 7:**
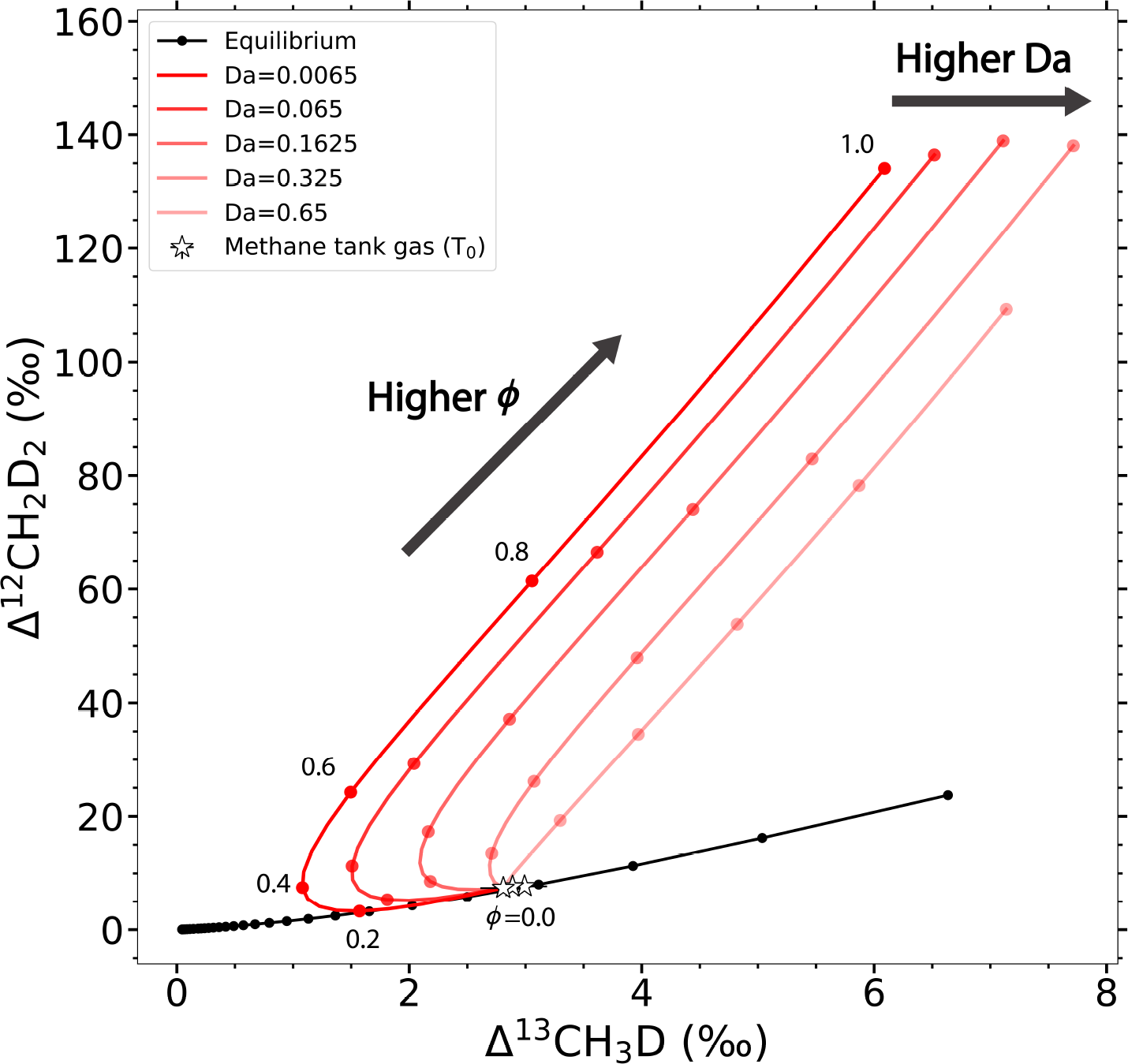
Steady-state clumped isotope signatures in the open-system model. Each curve represents one value of Da, and each point on a curve represents a specific *ϕ* (marked by the number by the point). The directions of increasing *ϕ* and Da are shown by the two arrows in the figure.

### 4.3. Implications

The results of this study reflect the sensitivity of the oxidation trajectory in clumped isotope phase space to net fractionation factors. Analogous results were obtained in some previous studies (Wang et al., 2016; Whitehill et al., 2017; Ono et al., 2021; Krause et al., 2022; Liu et al., 2023). We propose that variations in fractionation factors can arise in closed systems by changes in the relative magnitudes of cross-membrane methane transport and methane oxidation rate (i.e., Dahmköhler number, Da). When transport rate is high relative to oxidation rate, the oxidation trajectory resembles Rayleigh fractionation, and the net fractionation factors are close to the fractionation factors of methane oxidation. However, the net fractionation factors shift towards unity when the transport rate is about the same order of magnitude as the oxidation rate, causing the oxidation trajectory to deviate significantly from a Rayleigh fractionation trend (Figure 5). This effect is more prominent for Δ^12^CH_2_D_2_ as it reverses the sign of the trend (Figure 5, Figure 6). Therefore, the growth temperature (growth rate) does not alone control the clumped isotope ratios. Rather, temperature controls relative rates of oxidation and diffusion, which in turn determines the value for Da (eqn. 16) and thus isotope fractionation. The combined result of cross-membrane transport and methane oxidation determines the isotopologue effects on residual methane, requiring a distinction be made between experimentally-derived fractionation factors and the actual fractionation factors of aerobic oxidation. Systems with low methane (or oxygen) concentration and fast methane consumption will exhibit large offsets between the fractionation factors intrinsic to the oxidation reaction and those observed.

Oxidation trajectories in methane clumped isotope space in closed systems at high Da values resemble those in an open-system oxidation trending towards steady state (Haghnegahdar et al., 2017; Whitehill et al., 2017; Krause et al., 2022), both being characterized by very positive and increasing Δ^12^CH_2_D_2_ values. In our two-box model for closed systems (Figure 4), the cells behave like the representative elementary volumes in open-system models in previous studies (Wang et al., 2016; Krause et al., 2022), but with an important difference: open-system models typically assume methane is advected into the system with constant isotopologue ratios, while in the closed system, the isotopologue concentrations of methane transported into the cells are changing.

The recently reported clumped isotope compositions of atmospheric methane, which are strongly influenced by AeOM, show less positive Δ^12^CH_2_D_2_ values than expected from the models in which the source was assumed to be near isotopologue equilibrium (Haghnegahdar et al., 2023). This highlights the strong dependence of end-point isotopologue compositions on the initial composition. In our experiment, the initial methane tank gas is at thermodynamic equilibrium at a temperature of about 165 °C. If the initial methane isotopologue compositions begin with much lower Δ^12^CH_2_D_2_ values, in the microbial field, but follow the same oxidation trajectory as observed in this study, the resulting residual isotopologue compositions are expected to be near the equilibrium curve in mass-18 isotopologue space, with slightly positive Δ^12^CH_2_D_2_ values (Figure 8). The same argument applies to the open-system model. Under these circumstances, methane gas from a microbial origin but affected by AeOM will no longer show the tell-tale negative Δ^12^CH_2_D_2_ values that are typical for microbial methanogenesis (e.g., Young et al., 2017), and may instead exhibit positive Δ^12^CH_2_D_2_ values resembling either thermodynamic equilibrium or high Δ^12^CH_2_D_2_ values similar to atmospheric methane (Haghnegahdar et al., 2023).

**Figure 8:**
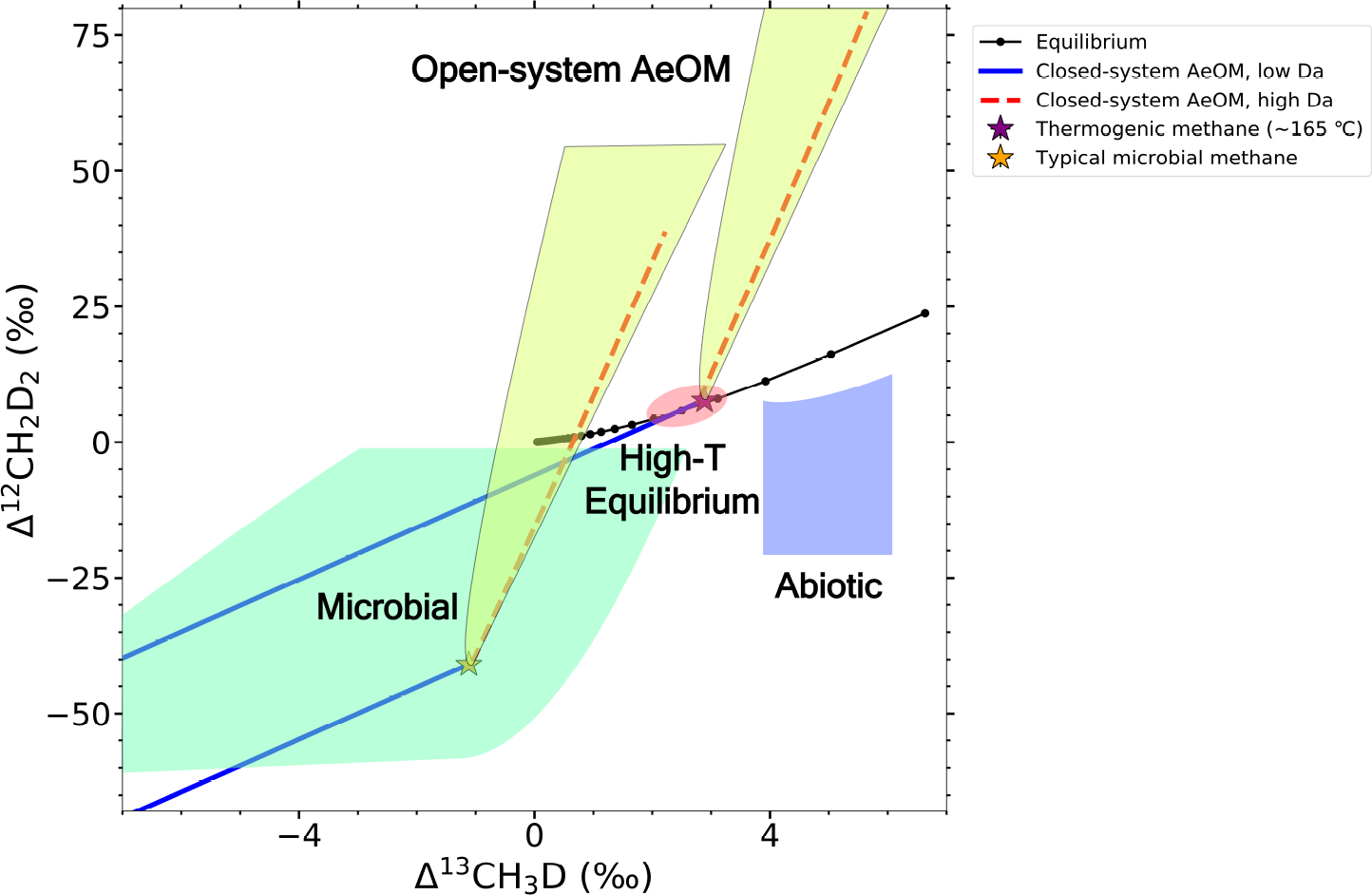
A summary of the potential effects of AeOM on the original clumped isotope signatures. The fields of clumped isotope values are adopted from Young et al. (2019). Stars indicate the starting clumped compositions of either thermogenic methane equilibrated at 165 °C, or disequilibrated microbial methane with Δ^13^CH_3_D = −1 ‰ and Δ^12^CH_2_D_2_ = −41 ‰. The red dashed lines and blue solid lines show the closed-system oxidation trajectories with high Da and low Da, respectively. The yellow shaded areas encompass the possible steady-state clumped isotope signatures of open-system AeOM.

Although the two-box model presented here fits the data very well, we are unable to quantify the variation of total methane transport with time in the experiments. Future studies with more accurate quantification of methane uptake and oxidation (e.g. using isotopically-labeled methane) are necessary to test the model.

Additionally, our model only accounts for total methane oxidation and transport, but cell-specific methane transport and methane oxidation rates may also influence the clumped isotope signatures, leaving considerable room for further study.

## 5. Conclusions

The growth rate effect on the isotope signatures (*δ*^13^C, *δ*D, Δ^13^CH_3_D and Δ^12^CH_2_D_2_) of residual methane left over from closed-system aerobic methane oxidation were investigated experimentally. Results show that the relative magnitudes of methane oxidation and cross-membrane methane transport, characterized by a Damköhler number, may influence the net isotopologue fractionation of the residual methane. We find that the variations in isotope fractionation observed in the experiments are not related to the switch of the active methane monooxygenases in the system nor to the hydrogen isotope exchange with water. Rather, our twobox model consisting of cellular methane, headspace methane, cross-membrane methane exchange, and intracellular methane oxidation naturally accounts for shifts in fractionation factors in the experiments as growth progresses. The model indicates that the Damköhler number strongly influences clumped isotope signatures in closed systems, as it can completely change the sign of Δ^12^CH_2_D_2_. In contrast, the general trend of oxidation is not strongly affected by Da in open systems, although the trends for open-system oxidation and closed-system oxidation at low Damköhler number are similar.

These results underscore that it is crucial to consider the potential influence of aerobic methane oxidizing microorganisms when using bulk and clumped isotope compositions to diagnose the provenance of methane gases, especially in the environments where methane or oxygen are not abundant. Further quantitative studies on the relation between methane transport and oxidation rates are required to validate the model developed above. This study provides insights into the key factors that control clumped isotope fractionation in closed-system methane oxidation, and along with previous studies, contribute to the development of methane clumped isotope as a tracer of methane cycling.

## Acknowledgements

This research was supported by funding from NASA grant 80NSSC21K0477 (EDY, WDL) and the Simons Foundation Award #623881 (WDL). We thank Whitney Thomas, Carolynn Harris and Jiarui Liu for the assistance on the lab experiments and thank Jeemin Rhim for the insightful discussions on the project.

## Data Availability

Data and codes are available through GitLab at https://git.dartmouth.edu/leavitt_lab/aerobic_methanotroph and figshare at https://figshare.com/projects/Li_etal_2024_AeOM/184504.

## Appendix A. Modeling

This includes the ordinary differential equation set and the parameters used in the two-box model.

### Appendix A.1. The ordinary differential equation set for the two-box model

This section shows the ordinary differential equation set that is used to calculate the evolution of methane isotopologues with time. The “o” and “i” in the subscripts refer to the outside and inside boxes, and the “T” in the subscripts refers to total methane. The *k*’s in the eqn. A.6-A.18 are the rate constants controlled by *α* and a tunable coefficient *K*_oxidize_ (Table 3 in the main text).

For the box outside the cell:

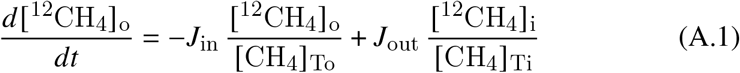

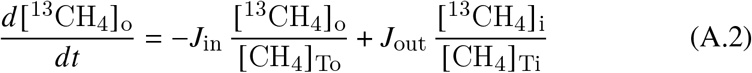

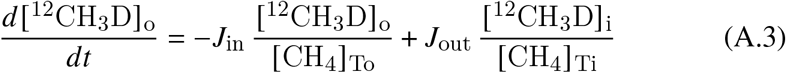

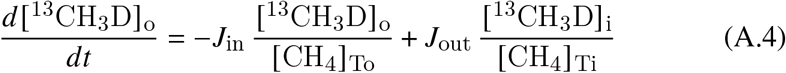

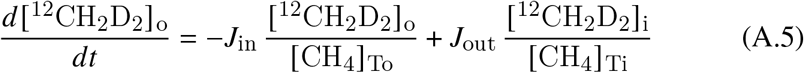

For the box inside the cell:

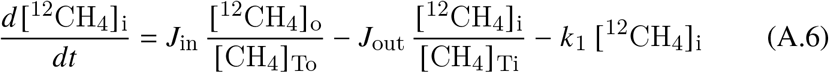

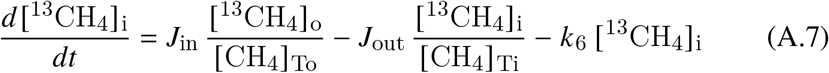

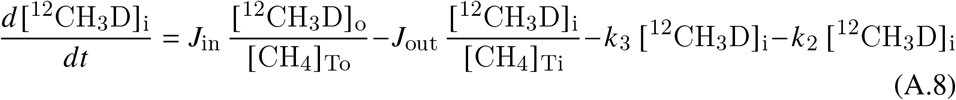

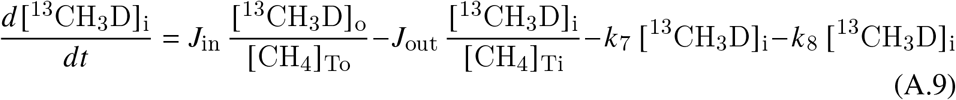

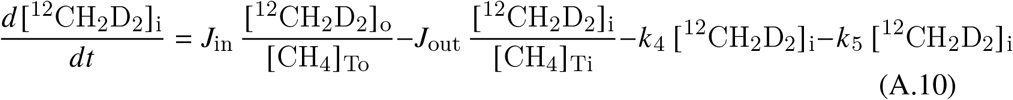

Other radicals inside the cell (not really considered in this model as the reactions are irreversible):

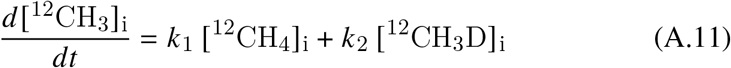

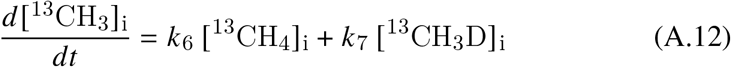

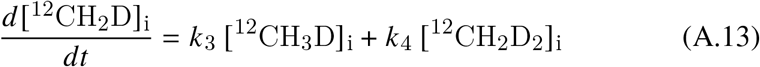

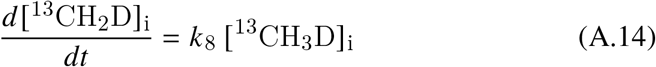

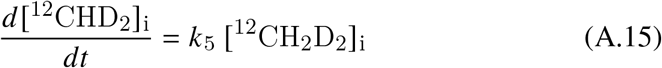

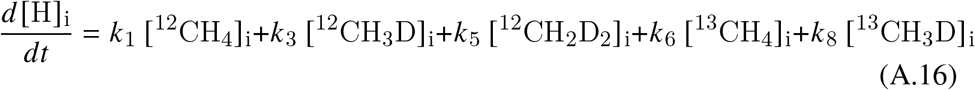

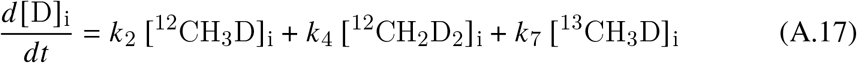

Total methane oxidation rate:

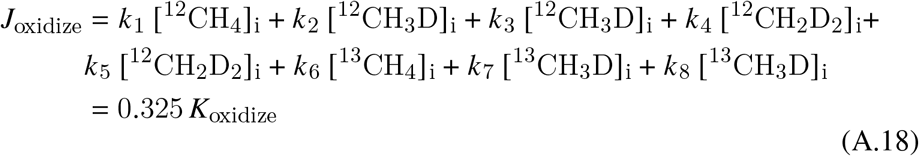

### Appendix A.2. Parameters in the model

Total methane: 650 *μ*mol;

Total reaction time: 28.0 (37 °C), 195 (27 °C), 508 (21 °C) hrs;

Total time steps: 10^5^.

## Appendix B. Supplementary figures

This contains Supplemental Figures B.9 to B.11 that show: growth curves and headspace methane concentrations (Figure B.9), dissolved copper concentrations in the media (Figure B.10), and the fitted curves for log(OD), Da and *J*_*in*_, *J*_*oxidize*_ with time (Figure B.11).

**Figure B_9:**
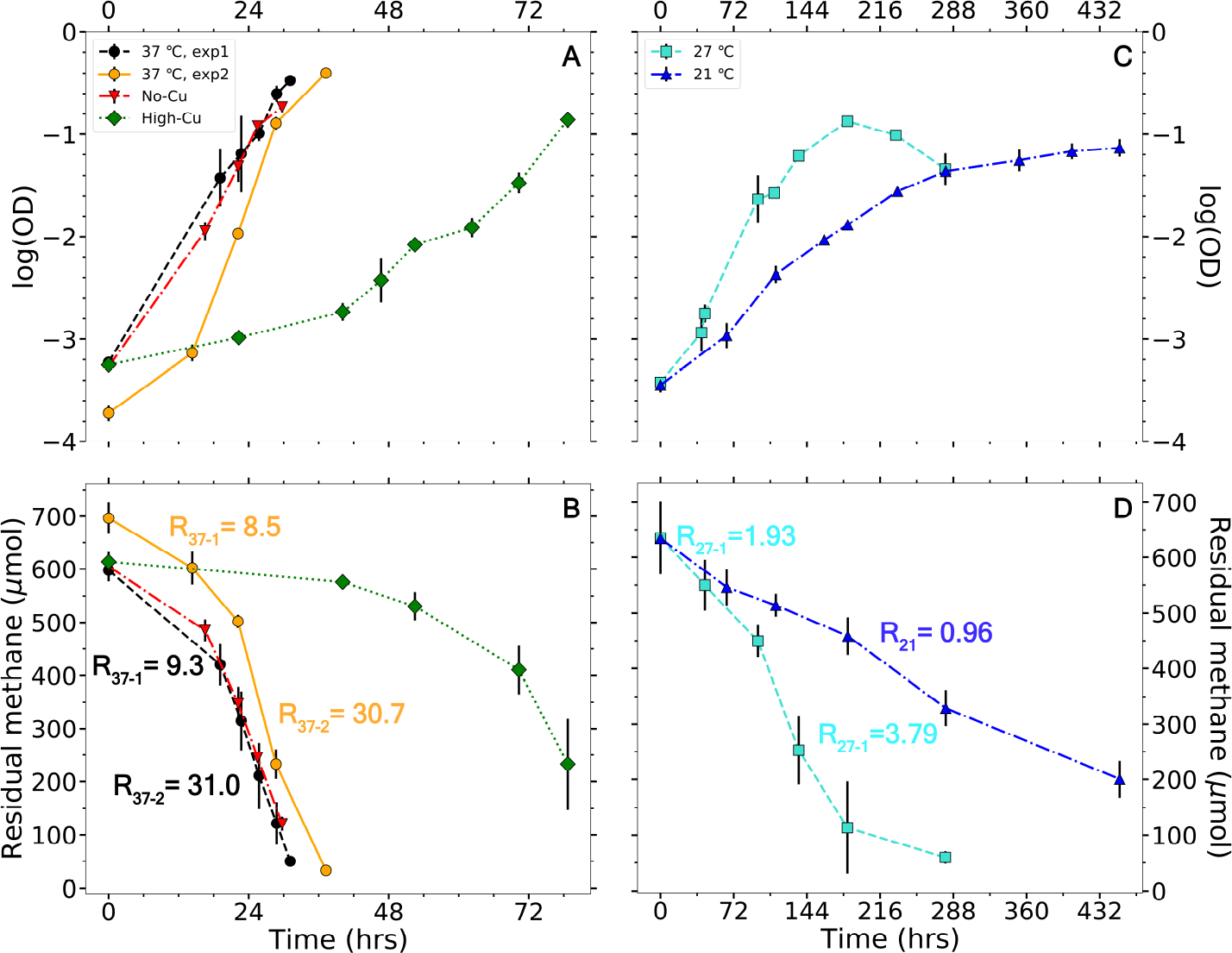
The growth curves and residual headspace methane amounts of the experiments. Panel A and B show the growth curves and headspace methane amounts under 37 °C and with different copper concentrations. Panel C and D show the growth curves and headspace methane amounts under 27 and 21 °C. Change of total methane oxidation rates in the 37 and 27 °C experiments can be observed from the change of the slopes in panel B and D. The total methane oxidation rates (in *μ*mol/hr) in the differential temperature experiments are shown in panel B and D by the curves.

**Figure B_10:**
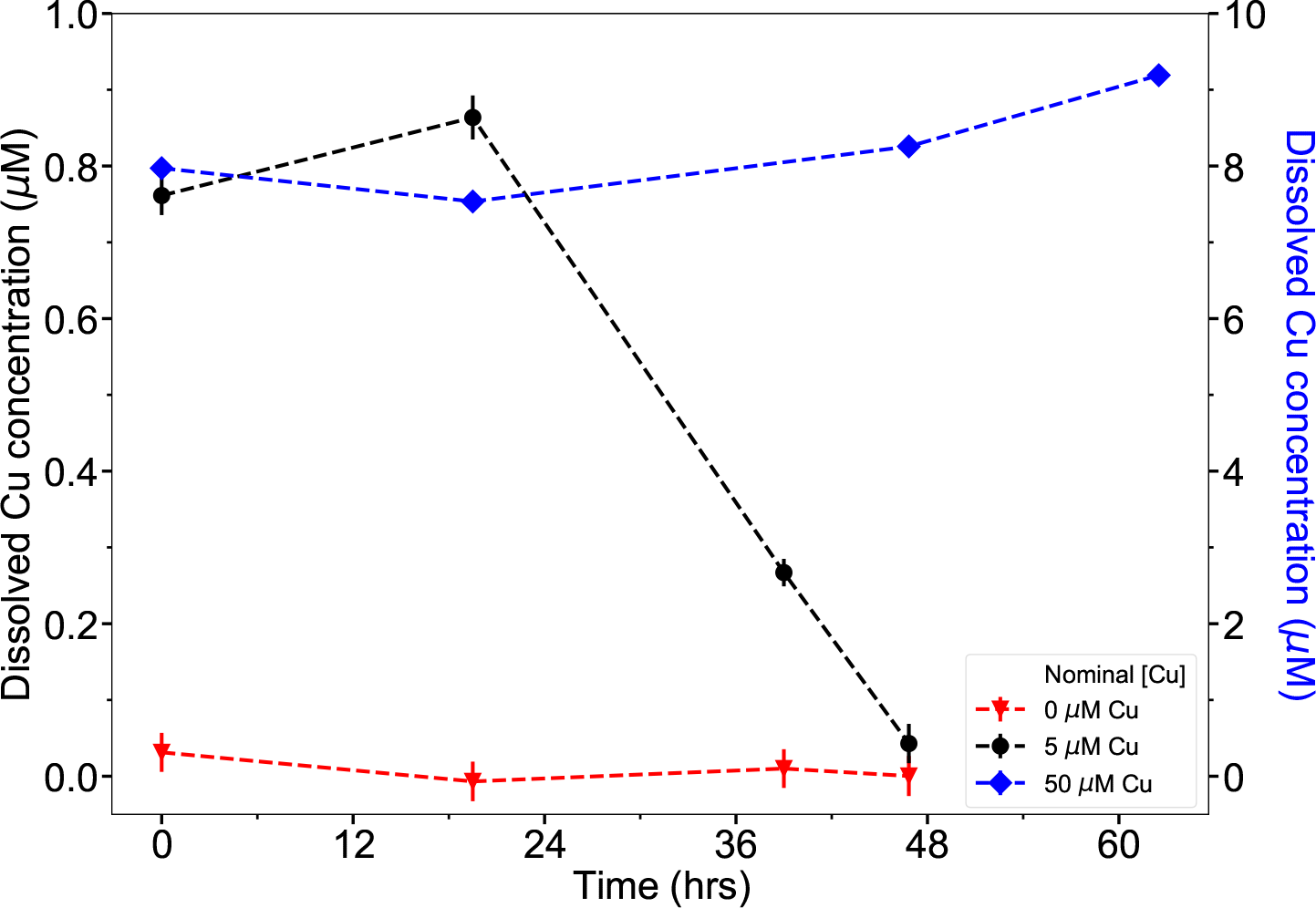
Dissolved copper concentrations ([Cu^2+^] _aq_) in the media under 37 °C with 0, 5 and 50 *μ*M Cu added. These are the nominated copper concentrations, but the actual dissolved copper concentrations are lower due to the precipitation of copper salts. Since [Cu^2+^] _aq_ in the experiment with 50 *μ*M Cu is an order of magnitude higher than the experiment with 5 *μ*M Cu, its values are shown on the y-axis at the right.

**Figure B_11:**
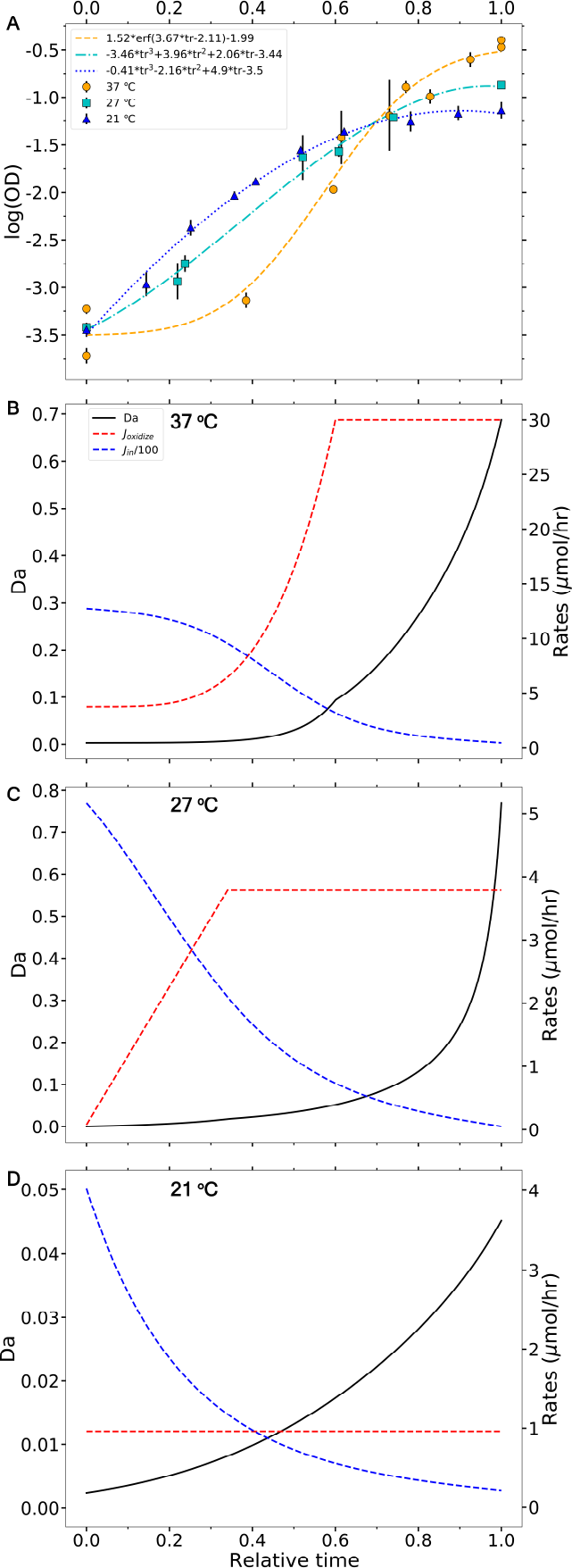
Fitted curves of log(OD) (panel A) and temporal trend of Da numbers, *J*_in_ and *J*_oxidize_ (panel B-D) in the models for 37, 27 and 21 °C experiments. Since the total time in the model is different between three experiments, for consistency, the x-axes show the time relative to the total modeled time (i.e. relative time, tr) in each experiment. tr = 0 and 1 mark the beginning and the end of the model, respectively. The log(OD) at 37 °C is fitted with an error function (erf) and the other two are fitted with third-order polynomial functions. The left axes in panel B-D are the Da numbers and the right axes are the rates *J*_in_ and *J*_oxidize_. The values of *J*_in_ are divided by 100 to scale with *J*_oxidize_.

## Appendix C. Supplementary tables

This includes Supplemental Tables C.4 and C.5 for dissolved copper concentrations and water *δ*D, respectively.

**Table C_4:**
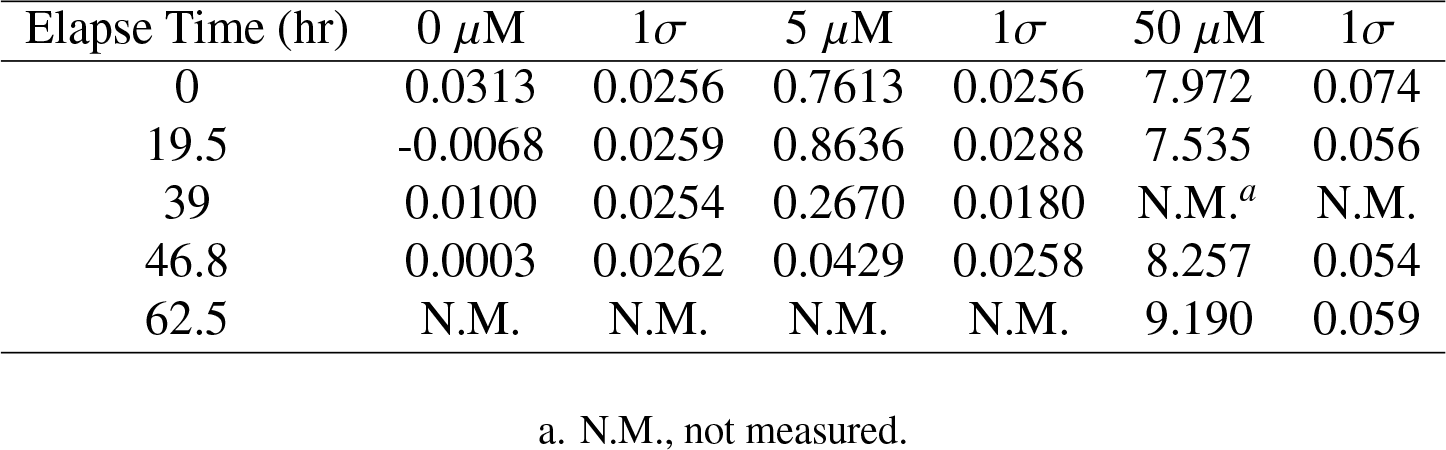
Measured dissolved copper concentrations in the media with time.

**Table C_5:**
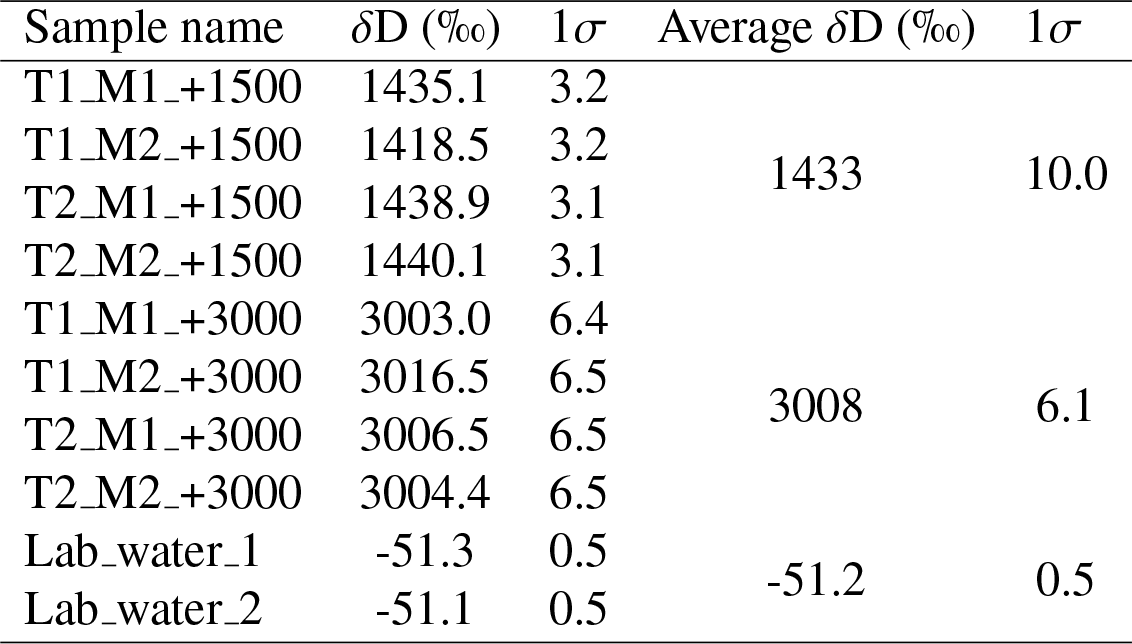
Measured water *δ*D in non-spiked and D-spiked experiments. Four samples are measured for each D-spike, with the uncertainties of dilution and isotopic measurement propagated into the 1*σ* error of each sample. Two lab water samples are measured to represent water *δ*D in the non-spiked experiments. The average *δ*D values are reported as the final results. Errors of the average values are reported as the 1 standard deviation of the four measurements.

